# Cost-information trade-offs partially rationalize the feed-forward loop census

**DOI:** 10.64898/2026.11.25.769197

**Authors:** Rafael Velásquez, Juan Manuel Pedraza

## Abstract

Gene transcription regulatory networks let cells encode an internal representation of their environment as patterns of protein expression. Among the thirteen connected directed subgraphs on three nodes, the feed-forward loop (FFL) is the one consistently over-represented in transcriptional networks relative to randomizations that preserve the node degrees, yet its eight structural variants occur very unequally: in *Escherichia coli* and *Saccharomyces cerevisiae* the C1 and I1 variants dominate and the other six are rare. We ask whether the trade-off between cost and information in these architectures can rationalize this asymmetry. Each variant is treated as a noisy channel from the environment to the output expression level, with a channel capacity obtained from the Lya-punov equation of the linearized chemical Langevin dynamics and a protein burden from the mean of the regulated proteins at steady state. The analytical capacity agrees with mutual information estimated from Gillespie simulations across families, gates, and Hill coefficients. We compare the eight variants on the Pareto frontier of the cost-information plane under four information measures and ten promoter logics. Two results follow. First, only C1, C3, I1, and I3 are ever Pareto winners within their family, while C2, C4, I2, and I4 never win, consistent with their rarity; C1 robustly dominates the coherent family, in agreement with the census. Second, no measure separates I1 from I3 by more than a few percentage points, and the measures disagree on which is ahead, so cost and information treat the two as nearly equivalent. In the census, by contrast, I1 outnumbers I3 by 21:1 in *S. cerevisiae*. Cost and information thus account for the exclusion of four variants and for the dominance of C1, but not for the dominance of I1, whose separation from I3 may require a measure of response timing, which our measures of the environmental level do not probe.

## 1. INTRODUCTION

Gene transcription regulatory networks (GTRNs) underlie essentially every physiological program of a cell, from sensory transduction to developmental decisions [1]. Their outputs are gene products, and one subset of these products, the transcription factors (TFs), closes the regulatory loop by binding DNA and modulating the transcription of other genes [1, 54]. TFs are conventionally divided into activators, which up-regulate their targets, and repressors, which down-regulate them; the combined action of many TFs on many promoters defines a directed graph whose nodes are genes and whose edges are regulatory interactions. Although regulation also operates post-transcriptionally and post-translationally [55], the transcriptional layer alone contains a rich repertoire of patterns (cascades, fan-ins, fan-outs, mutual pairs, feedback and feed-forward loops) that has been mapped for organisms from *Escherichia coli* and *Saccharomyces cerevisiae* to mouse and human [1–4] [Fig. 1(a)].

**FIG 1.**
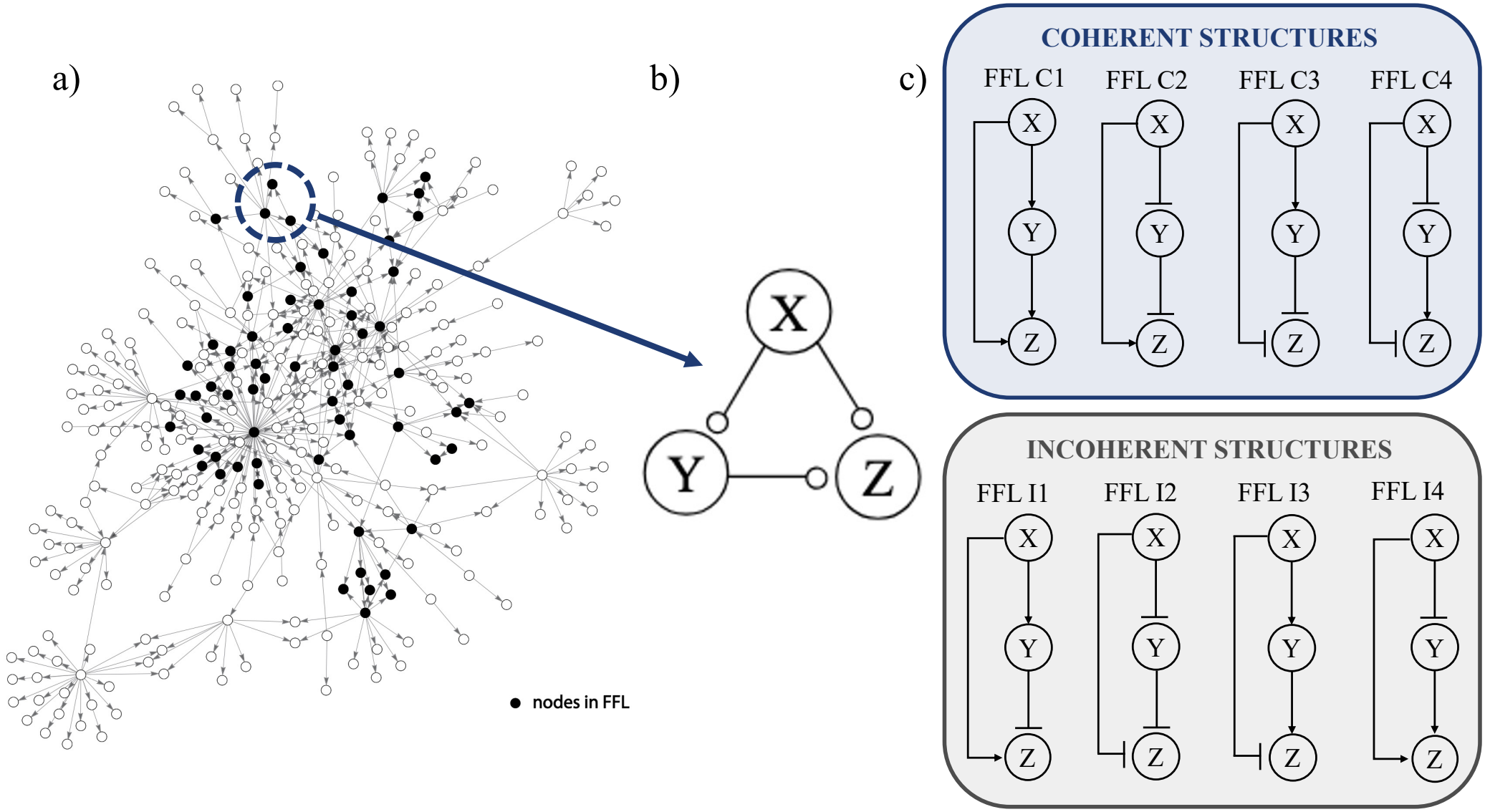
The biological context of the present study. **(a)** A *schematic* representation (not adapted from any particular data set) of a gene transcription regulatory network: nodes are genes, directed edges are transcriptional regulations, and a small number of nodes (the hubs, in lighter blue) act as master regulators of many targets. Among all three-node subgraphs of such networks, only the *feed-forward loop* (FFL) is statistically over-represented relative to randomized graphs with the same in– and out-degree distributions in the systems analyzed in Ref. [2, 18]; one such FFL is highlighted in red. **(b)** Schematic of an isolated FFL with input *X*, intermediate *Y*, and output *Z*, and an environmental input that modulates the transcription rate *K*_*X*_ of the upstream gene. **(c)** The eight structural variants of the FFL obtained by assigning each of the three regulatory edges a sign (activation, →, or repression, ⊣). Variants are coherent (C1–C4, green) when the sign of the direct edge *X* → *Z* coincides with the product of the signs of the indirect path *X* → *Y* → *Z*, and incoherent (I1–I4, pink) otherwise. Despite their structural symmetry, the natural census of these eight variants is highly asymmetric, with C1 and I1 dominating the empirical abundance reported in Ref. [21].

A central function of these networks is to let the cell sense and respond to its environment [1]. Lacking a centralized controller, the cell encodes its decisions as patterns of expression: intracellular protein concentrations form an internal representation of the external state, and the regulatory network updates that representation as upstream signals fluctuate [50, 51]. Transcriptional regulation can therefore be cast as information transmission [5, 51], with a noisy environmental signal as input, the expression level of an effector protein as output, and the network as the noisy channel [6, 7] between them; the physical limits of this transmission and the information-theoretic analysis of regulatory circuits have been reviewed extensively [50, 52]. The figure of merit is the mutual information between environment and output, and a substantial body of work indicates that biological signaling networks operate close to information-theoretic optima [8–1 Information transmission has been analyzed specifically for feed-forward loops, both for the optimization of their input-output relation [47] and for the frequency-dependent transmission that distinguishes coherent from incoherent variants [48, 49]. At the same time, fundamental limits on noise suppression [15, 52] and the energetic costs of biochemical computation [16, 17] bound the resources required for a given fidelity, so optimality in transcriptional networks must be discussed with information and cost on the same footing.

A surprising organizational principle of these networks was identified by Milo, Alon, and collaborators [2, 18]: among the thirteen connected three-node directed sub-graphs, only the *feed-forward loop* (FFL) is statistically over-represented in transcriptional networks relative to randomized graphs that preserve the in– and out-degree distributions [3]. The strength of this enrichment depends on the network definition and on the randomization ensemble, but the FFL is consistently recovered as a motif in the organisms that have been studied carefully (*E. coli, S. cerevisiae*, mouse, and human). It has therefore been proposed as an elementary computational building block favored by evolution, and the notion has since been extended to larger subgraphs and motif clusters [19, 20].

Even within the FFL family, however, the natural census is highly asymmetric. Each of the three regulatory edges can be an activation or a repression, giving eight structural variants that split into four *coherent* (C1–C4) and four *incoherent* (I1–I4) FFLs [21] (Sec. II and Fig. 1). In the *E. coli* and *S. cerevisiae* transcriptional networks, C1 (three activations) dominates the coherent family and I1 (activation *X*→ *Y*, repression *Y*⊣ *Z*, activation *X*→ *Z*) dominates the incoherent family [21]. Why is the census so asymmetric, and which selection pressures are responsible?

Functional arguments have been advanced for individual variants, including persistence detection by C1 [22, 23], response acceleration by I1 [24], fold-change detection [25], and pulse generation [24]. These analyses typically rely on deterministic, mean-field dynamics and do not compare the frequencies of all eight variants on a common quantitative footing. A complementary line of work embedded FFL variants in environmental-selection scenarios and showed that C1 is favored under fluctuating inputs that require persistence filtering [26]; this establishes that fitness, and not topology alone, shapes FFL abundance, but it does not identify which quantitative property of the dynamics is optimized. Here we adopt an information-theoretic point of view. We treat each FFL as a noisy channel that transmits information about the environment to the output *Z* and quantify its performance by the mutual information *I*(*E*; *Z*), where *E* is encoded in the input transcription rate of the upstream gene *X* [6, 7]. Because *X* itself fluctuates, the relevant question is not how well *Z* tracks *X* but how well the whole network reports the environmental state. Sustaining this transmission requires the production of intermediate and output proteins and therefore consumes metabolic resources [27–29], so we compare variants through the Pareto frontier of the costinformation plane rather than through a single benefit-cost ratio. Relative to previous work, we place information and cost on an equal quantitative footing for all eight variants simultaneously, evaluate them with four information measures across ten gate logics, and support the comparison with bootstrap confidence intervals, paired tests with multiple-comparison correction, and analyti-cal/numerical cross-validation.

Our analysis combines two approaches. Analytically, we linearize the chemical Langevin equations around the deterministic steady state and solve the Lyapunov equation for the conditional output variance 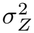 (*E*); together with the local gain *g*(*E*) = *d* ⟨*Z*⟩ */dE*, this yields the Brunel-Nadal-Laughlin channel capacity in the small-noise limit [8, 42, 43]. Numerically, we run Gillespie simulations [38] and estimate *I*(*E*; *Z*) from discrete environmental states, both at steady state and after step changes of the input. The first approach gives a capacity (the mutual information maximized over input distributions) and the second the mutual information achieved under a fixed prior; we report both and use their agreement as an internal consistency check (Sec. IV). We also consider ten two-input gate logics at the *Z* promoter rather than only AND and OR, and we test the robustness of the within-family ranking to alternative cost specifications and to reweighting of the kinetic-parameter prior. The framework identifies the dominance of C1 within the coherent family but treats I1 and I3 as nearly equivalent: no information measure separates the two by more than a few percentage points, and the measures disagree on which is ahead. The empirical dominance of I1, which the census favors over I3 by more than 5:1, therefore cannot be attributed to cost and information alone.

The paper is organized as follows. Section II reviews the eight FFL variants and the gate logics at the *Z* promoter. Section III introduces the stochastic model. Section IV derives the small-noise channel capacity *C* and validates it against Gillespie estimates of *I*(*E*; *Z*). Section V defines the protein burden and the Pareto analysis and tests their robustness to the cost specification and to the parameter prior. Section VI presents the capacity landscape, the stratified Pareto shares under four information measures, the gate-simplex analysis, and the unstratified comparison, and Sec. VII discusses the implications for the FFL census.

## II.THE FEED-FORWARD LOOP AND ITS VARIANTS

A feed-forward loop is a directed subgraph on three nodes *X, Y, Z* with the following edges: *X*→ *Y*,

*X* → *Z*, and *Y* → *Z*. Each of the three edges can be either an activation (+) or a repression (−), giving 2^3^ = 8 structural variants. We label them following the standard nomenclature of Mangan and Alon [21].

A variant is called *coherent* if the sign of the direct path *X*→ *Z* coincides with the sign of the indirect path *X*→ *Y* →*Z* (the latter being the product of the signs of *X*→ *Y* and *Y*→ *Z*). Otherwise, the variant is called *incoherent*. The four coherent variants are denoted C1, C2, C3, C4, and the four incoherent variants I1, I2, I3, I4 (Fig. 1).

In the language of signal processing, a coherent FFL is a circuit in which the direct and indirect paths reinforce each other, while an incoherent FFL is one in which they compete. Both architectures are useful, but in qualitatively different regimes: coherent loops act as persistence detectors and noise filters, while incoherent loops can generate pulses or accelerate responses.

Independently of the activation/repression pattern of the three edges, the target node *Z* integrates two regulatory inputs (*X* and *Y* ) and must therefore implement some combinatorial logic at its promoter. There are sixteen possible truth tables on a binary input pair; six of these are biologically trivial (constants and projections onto a single input) and the remaining ten genuinely combine both regulators. The two canonical limits are an AND gate (both regulators required) and an OR gate (either regulator sufficient), and previous information-theoretic treatments of the FFL have focused on these two cases. Here we consider all ten nontrivial input functions: AND, OR, and eight additional logics drawn from a thermodynamic occupancy model of the regulated promoter. Their explicit form is summarized in Sec. III B and derived in Appendix A.

## III. STOCHASTIC MODEL OF FFL DYNAMICS

We model each FFL as a system of six coupled chemical Langevin equations, one for the mRNA and one for the protein concentration of each of the three nodes. Denoting by *m*_i_ and *i* (*i* = *X, Y, Z*) the mRNA and protein levels respectively, the dynamics read levels respectively, the dynamics read

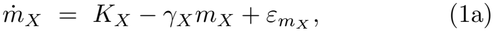

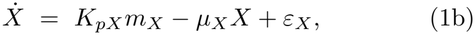

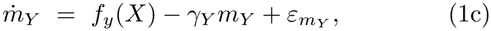

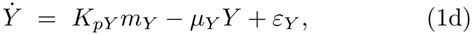

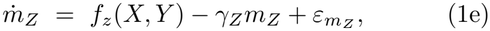

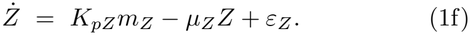

Here *K*_X_ is the constitutive transcription rate of *X, K*_pi_ is the translation rate of node *i, γ*_i_ is the mRNA degradation rate, and *µ*_i_ is the protein degradation/dilution rate. The functions *f*_y_(*X*) and *f*_z_(*X, Y* ) encode the regulatory logic of the loop and are detailed below. The terms 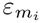, *ε*_i_ are independent Gaussian white-noise sources with zero mean and a variance set by the local rates of birth and death of each species, in the spirit of the linear-noise approximation [34].

The environmental state *E* is encoded in the constitutive transcription rate of the upstream gene, *E* ≡ log_10_ *K*_X_. Biological signal ranges are typically log-distributed, and this choice places the capacity calculation of Sec. IV in the framework of Tkačik and collaborators [8, 9], who also parametrize environmental states by the logarithm of an input concentration.

### A. Hill functions for single regulators

The regulation of *Y* by *X* depends on a single transcription factor and is described by a Hill function. For an activator,

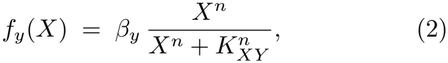

while for a repressor,

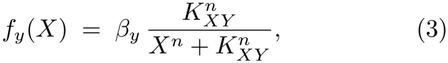

where *n* is the Hill coefficient and *K*_XY_ is the dissociation constant. We adopt the convention that Hill functions return values in [0, 1] (the regulator’s active-state occupancy), pre-multiplied by the maximal transcription rate *β*_y_. This convention will make the structure of the ten two-input functions *f*_z_(*X, Y* ) below transparent: each is built by applying a logic operation to the occupancies *g*_X_, *g*_Y_ ∈ [0, 1] of the two inputs, with the FFL’s signs determining whether the activator or repressor form of the Hill function is used for *X* and *Y* at the *Z* promoter.

### B. Ten two-input gate logics at *Z*

Defining

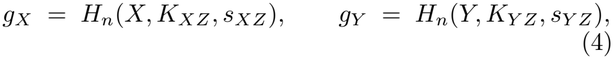

where *s*_XZ_, *s*_Y Z_ ∈ {+1, −1 } are the signs of the *X*→ *Z* and *Y*→ *Z* edges and *H*_n_ is the Hill activator form when *s* = +1 and the Hill repressor form when *s* = − 1, the two canonical Boolean limits read

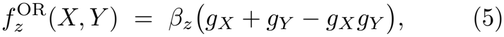

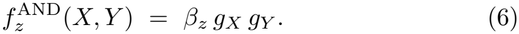

The remaining eight nontrivial logics, derived from a thermodynamic occupancy model of the regulated promoter (Appendix A), are[eq

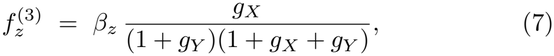

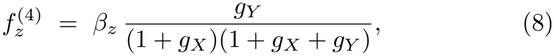

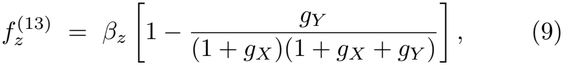

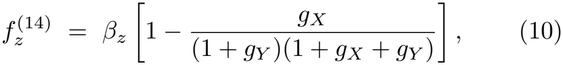

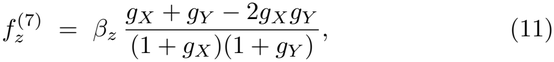

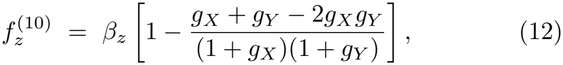

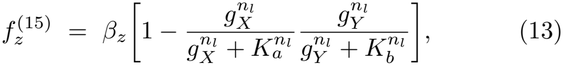

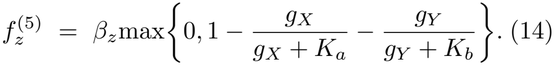

Because the edge signs already set the activator or repressor form of *g*_X_ and *g*_Y_, logics that invert an input (L3, L4, L5, L13, L14, L15) or respond non-monotonically to it (L7, L10) change the effective sign with which *X* or *Y* acts on *Z*; variant labels always refer to the edge signs of Sec. II. The numbering follows the underlying thermodynamic enumeration [45, 46]. Logics 3 and 4 are activator-modulated forms with the roles of *X* and *Y* exchanged, and logics 14 and 13 are their respective complements; logics 7 and 10 are XOR– and XNOR-like rational functions; logics 15 and 5 are NAND- and NOR-like Hill complements with internal cooperativity *n*_l_ and constants *K*_a_, *K*_b_, set to *K*_a_ = *K*_b_ = 0.5, *n*_l_ = 2 for L15 and to *K*_a_ = 0.5, *K*_b_ = 1, *n*_l_ = 1 for L5, within the regime in which the thermodynamic model has been characterized [46]. All analyses below use the AND and OR gates, except the gate-simplex analysis of Sec. VI F and App. C 4, which use all ten.

### C. Noise terms

Each Langevin equation in (1) carries an additive Gaussian white-noise term *ε* with ⟨*ε*(*t*)⟩ = 0 and ⟨*ε*(*t*)*ε*(*t*^′^ ) ⟩ = *D δ*(*t*−*t*^′^ ). The noise sources for different species are taken to be uncorrelated, ⟨*ε*_i_(*t*)*ε*_j_(*t*^′^)⟩ ∝ *δ*_ij_*δ*(*t t*^′^), since the elementary birth and death events of any two species are statistically independent. The diffusion constants *D* are fixed self-consistently in the linear-noise approximation by the steady state of Eq. (1), with diagonal entries *D*_ii_ = production_i_ + degradation_i_ summing the forward and backward fluxes affecting species *i* [34–36].

## IV.CHANNEL CAPACITY FROM THE LYAPUNOV EQUATION

The cell receives information about its environment *E* through the output *Z*, and the figure of merit for that channel is the mutual information *I*(*E*; *Z*). To make analytical progress we linearize Eq. (1) around the deterministic steady state at each environmental value *E* = log_10_ *K*_X_. Writing each species *ρ* ∈ {*m*_X_, *X, m*_Y_, *Y, m*_Z_, *Z* }as *ρ*(*t*) = ⟨ *ρ*⟩ (*E*) + *δρ*(*t*), with ⟨*ρ*⟩ (*E*) the *E*–dependent steady-state mean, and expanding the Hill and gate functions to first order in the fluctuations, we obtain

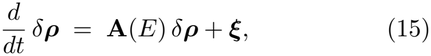

where *δ****ρ*** = (*δm*_X_, *δX, δm*_Y_, *δY, δm*_Z_, *δZ*)^⊤^, ***ξ*** collects the six noise terms, and **A**(*E*) is the Jacobian matrix evaluated at ⟨ ***ρ*** ⟩ (*E*). The Jacobian has the block-triangular structure

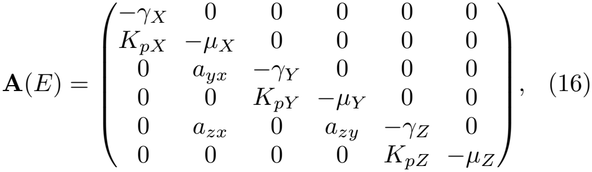

where *a*_yx_ = *∂*_X_ *f*_y_, *a*_zx_ = *∂*_X_ *f*_z_, and *a*_zy_ = *∂*_Y_ *f*_z_ are evaluated at the deterministic steady state; closed forms for the AND and OR gates are given in App. A 6. The block-triangular structure reflects the directed, acyclic topology of the FFL: *X* is regulated only by its own constitutive production, *Y* only by *X*, and *Z* by both *X* and *Y* .

The stationary covariance matrix **Σ**(*E*) of the fluctuations,

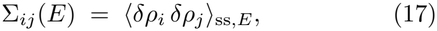

satisfies the Lyapunov equation [34, 37]

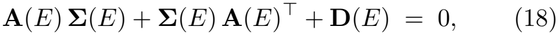

where **D**(*E*) = ⟨***ξ ξ***^⊤^⟩ is the diffusion matrix obtained from the chemical Langevin approximation. Equation (18) is a linear algebraic system for the symmetric entries of **Σ**(*E*) that we solve numerically using standard routines. The element of interest is the conditional variance of the output,

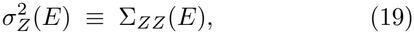

which sets the noise floor at environmental state *E*, together with the local gain

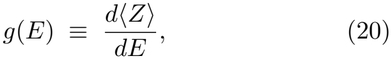

which sets how strongly the output depends on the environment near that operating point.

### A. Local Gaussian mutual information

For a Gaussian prior 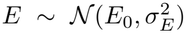 of width *σ*_E_ around an operating point *E*_0_, the linearized channel *Z* = ⟨*Z*⟩(*E*_0_) + *g*(*E*_0_)(*E* − *E*_0_) + *ξ* with 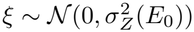 has mutual information

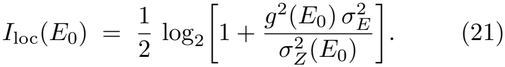

Equation (21) is the standard signal-to-noise form of the mutual information of a Gaussian channel [6, 7]. Its dependence on *E*_0_ shows where the output is informative: more bits are transmitted where the gain is large relative to the noise. The monotonic responses of coherent variants give single-peaked *I*_loc_(*E*) profiles, and the band-pass responses of incoherent variants give two-peaked ones (Sec. VI A).

### B. Channel capacity

Equation (21) assumes a fixed prior. The operationally relevant quantity for an evolved sensor may instead be the *channel capacity*, the maximal mutual information over all input distributions [6, 8]. For a small-noise Gaussian channel with input-dependent gain *g*(*E*) and noise *σ*_Z_(*E*), the capacity takes the closed form [8, 42, 43]

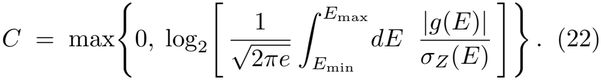

The integrand is the square root of the local Fisher information 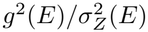, so the integral counts the distinguishable output levels over the operational range of *E*. The outer max{0, ·} sets *C* = 0 when the integral falls below 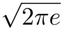, where the small-noise expression would become negative. This floor is reached: it clips *C* to zero for about a quarter of the parameter samples, almost all of them low-information regimes where the Gillespie estimate is also small (Sec. IV D), and a robustness check confirms that discarding these samples does not change the within-family rankings (Sec. VI C). At the representative point of Sec. VI A the lowest incoherent capacities (0.10 and 0.36 bits for I2 and I3) sit just above the floor. The capacity-achieving input distribution is *p*^⋆^(*E*) ∝ |*g*(*E*) | */σ*_Z_(*E*), which weights the environmental states where the channel discriminates best.

Our two information measures are therefore distinct. The Lyapunov approach gives the capacity *C*, achieved by *p*^⋆^(*E*); the Gillespie approach gives the mutual information *Î*_SSA_(*E*; *Z*) under a fixed uniform prior over eight environmental states (Sec. IV C). For an evolved sensor it is not obvious which is the better figure of merit: the capacity describes the best the architecture could do if the input statistics were matched to the channel, whereas *Î*_SSA_ describes performance at one particular prior. We report both and treat their agreement (Sec. IV D) as evidence that our conclusions do not depend on this choice.

Equation (22) assumes that the mapping *E* ↦⟨*Z*⟩ (*E*) is locally one-to-one, so that the local Fisher information captures the distinguishability of nearby environmental states. This holds globally for coherent variants, whose dose-responses are monotonic. For incoherent variants the dose-response is non-monotonic, and two states on opposite sides of its maximum can produce the same mean output. Because the integrand |*g*(*E*) |treats each branch separately, the formula may overestimate discrim-inability by ignoring this branch degeneracy.

The integrand vanishes at the maximum itself, where *g*(*E*) = 0, but this does not remove the confusion between states on opposite branches. The decisive test is therefore a comparison with the Gillespie estimator *Î*_SSA_(*E*; *Z*), which makes no monotonicity assumption and includes any loss from branch degeneracy that is resolvable with eight environmental states and eight quantile bins of *P* (*Z* | *E*). Section IV D shows that the two estimators are strongly correlated in the incoherent sub-groups, only slightly less than in the coherent ones, and that their within-family information rankings are identical or differ by one adjacent transposition.

Eight states and eight bins might, however, be too coarse to resolve the degeneracy, in which case the agreement could reflect a shared blindness rather than mutual confirmation. We therefore recomputed the mutual information for representative parameter samples of I1 and I3 under OR gating at *n* = 2, one of the subgroups in which the stationary Pareto shares favor I3 (Sec. VI B), with three estimators of increasing resolution: the baseline plug-in estimator (eight states, eight bins), a finer plug-in estimator (forty states, thirty-two bins), and the binning-free Kraskov-Stögbauer-Grassberger (KSG) *k*– nearest-neighbor estimator [40, 41] on the fine grid. The three agree closely, both in the information carried by each variant and in the difference *I*_I1_ − *I*_I3_ (− 0.23, 0.26, and −0.27 bits, respectively), so the coarse binning does not hide structure. This check concerns the estimator only. None of our conclusions rests on fine structure of the incoherent response: the dominance of C1 concerns the monotonic coherent regime, where no degeneracy arises, the four-variant reduction holds under all four information measures, and the I3 lead over I1 under OR is also returned by the analytical capacity, which involves no binning.

### C.Numerical estimation from stochastic simulations

The Lyapunov calculation relies on linearization and on the Gaussian approximation. To test these, we also estimate *I*(*E*; *Z*) from Gillespie simulations [38] of the master equation of the reactions underlying Eq. (1). We use a discrete uniform prior over eight values 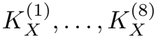 log-spaced in [1, 8], simulate replicate trajectories to steady state for each value, bin the output *Z*, and apply the plug-in estimator with Miller-Madow bias correction [39]. The same pipeline gives the transient mutual information of Sec. VI D.

### D.Quantitative validation of the small-noise approximation

To test quantitatively whether the small-noise approximation or the choice of prior affects the comparison, we evaluated *C* and *Î*_SSA_(*E*; *Z*) at the same *N* = 1000 log-uniform parameter samples in each (family, gate, *n*) sub-group (Fig. 6 in App. C 3). Over all 160 000 paired values (ten gates, two Hill coefficients, two families, four variants, and 1000 samples), the Pearson correlation between the two estimators is 0.82 across the pooled set and lies in [0.80, 0.95] within the individual subgroups. The residuals *C* −1.15 *Î*_SSA_ have mean −0.16, median −0.17 bits, and standard deviation 0.33 bits. They are approximately symmetric and show no systematic curvature against ⟨*Z*⟩ .

A correlation over this wide dynamic range certifies that the two estimators agree on which circuits transmit much or little information, not that either resolves a difference of a few tenths of a bit between two variants. The comparisons that carry our conclusions, however, are paired: I1 and I3 (or any two variants) are evaluated at the same parameter sample and through the same pipeline, so estimator biases that depend on the parameters or on the output scale enter both variants and cancel in the per-sample difference *D*_k_. The relevant resolution is therefore not the scatter of the absolute estimate but that of the paired difference, which is smaller; the bootstrap intervals on the shares (Table III) quantify it directly.

The capacity floor of Eq. (22) clips *C* to zero for 24.9% of the paired values. These clipped samples are concen-trated at low information: only 3607 of them (about 2% of all pairs) have *Î*_SSA_ *>* 0.5 bits, and the largest SSA estimate among clipped samples is 1.35 bits. The clipping therefore acts almost exclusively where the simulation also reports little information, so it removes few informative points; its effect on the rankings is quantified in Sec. VI C.

The unclipped aggregate slope of the linear fit is 1.11, so the capacity exceeds the SSA estimate by about 10% where both are nonzero. Two effects act in this direction: the capacity uses the optimal continuous prior *p*^⋆^(*E*), which extracts more information than the uniform eight-state prior, and it is integrated over *K*_X_ ∈ [0.5, 10] (App. B), whereas the SSA prior covers only *K*_X_ ∈ [1, 8]. Neither is an approximation error; both leave the within-family orderings unchanged, since they apply equally to all variants of a subgroup.

Finally, the within-family ranking of variants by mean information is identical under the two estimators in all four coherent subgroups (*ρ* = 1.00) and in one incoherent subgroup, and differs by a single adjacent transposition in the other three incoherent subgroups (*ρ* = 0.80; Table II). Because the two estimators use different priors, this concordance also shows that the rankings are not an artifact of the prior used in either pipeline.

## V.METABOLIC COST AND PARETO FRAMEWORK

Transmitting information is not free: maintaining in-termediate and output proteins consumes nucleotides, amino acids, and ATP and occupies ribosomes and RNA polymerases. This expression burden is measurable. Increasing the expression of a single unneeded operon in *E*.*coli* reduces growth rate, and the expression level evolves toward an optimum that balances burden against benefit [27–29]. Since protein synthesis dominates the cost of expressing a typical gene, we use the steady-state mean of the regulated proteins as a *protein-burden proxy*,

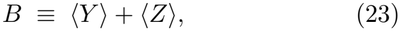

averaged over the operational range of *E*. Throughout the paper *B* denotes this burden and *C* the channel capacity. The input ⟨*X*⟩ is omitted because it is shared by all eight variants and drops out of any comparison between them.

A detailed biophysical cost would depend on protein length, amino-acid composition, codon usage, and turnover, none of which are equal for *Y* and *Z* in real systems. We use Eq. (23) for three reasons. First, titration and proteome-wide expression experiments show that growth rate decreases monotonically, and over the relevant range approximately linearly, with the burden of unneeded protein in *E. coli* [27] and *S. cerevisiae* [30], and translation-related contributions to this burden have been quantified in *E. coli* [31]. Second, length– and codon-related differences between *Y* and *Z* act as approximately constant factors on the two terms of Eq. (23), common to all variants; they reweight the two terms rather than reorder variants arbitrarily, and we test this reweighting below. Third, the within-family ranking is survives the linear, nonlinear, and single-species alternatives to Eq. (23) (Sec. V C).

At given parameters and gate, each variant occupies a point (*B, I*) in the cost-information plane. Rather than collapse this onto a benefit-cost ratio, which requires an arbitrary exchange rate between bits and proteins, we use the *Pareto frontier* : a variant is Pareto-optimal within a comparison set if no other variant in the set transmits at least as much information at no greater cost, with at least one of the two strictly better. The frontier is unchanged by any strictly increasing transformation of either axis [44].

### A. Pareto share and stratification

The frontier depends on the kinetic parameters of each realization. We therefore sample *N* parameter sets log-uniformly (App. B) and compute the frontier within a comparison set at each sample. If *k* variants lie on the frontier at a given sample, each receives a credit 1*/k*, and the *Pareto share* of a variant is its credit averaged over samples. Shares within a comparison set sum to unity and can be read as the fraction of frontier “mass” carried by each variant. A variant can carry a substantial share while rarely being alone on the frontier.

By default we compute the frontier separately for the coherent and incoherent families at fixed gate and Hill coefficient *n*, so that each comparison contains four variants (*stratified* analysis). In the *unstratified* control, all eight variants compete (Sec. VI G). Stratification by family has a biological motivation: coherent and incoherent FFLs implement different functions, persistence detection with monotonic dose-responses versus pulse generation with band-pass dose-responses [21, 24], so selection plausibly compares architectures within a functional class, and the census of Ref. [21] that we compare against is itself organized by family.

### B. Robustness to gate uncertainty

Which two-input logic a given promoter implements is generally unknown. We therefore treat the gate as a methodological variable and consider convex mixtures of the ten gates of Sec. III B. For a weight vector **w** on the nine-dimensional probability simplex Δ^9^, the gate-mixed Pareto share of a variant is *F*_w_(FFL) = ∑ _*l*_ *w*_*l*_*P*_*l*_(FFL), where *P*_*l*_ is its Pareto share under gate *l*. At each **w**, the within-family *sole winner* is the variant with the largest *F*_w_, and the *sole-winner mass* of a variant is the fraction of Δ^9^, under the uniform Dirichlet measure, on which it is the winner, estimated by Monte Carlo sampling of **w**. The mixtures interpolate between gate behaviors at the level of Pareto-share statistics; we do not claim that every point of Δ^9^ corresponds to a physically realizable promoter. The analysis therefore measures how broadly across regulatory logics a conclusion holds, not the underlying biochemistry.

### C. Robustness to alternative cost specifications

Equation (23) is one of several plausible cost quantifications. Using the same *N* = 1000 parameter samples, we tested three families of alternatives (Table I): linearly weighted costs *B* = *α*⟨*Y* ⟩ + *β*⟨*Z*⟩ with *α/β* ∈ [1*/*3, 3]; four nonlinear costs, 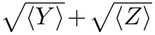, ⟨*Y* ⟩ ^2^ + ⟨*Z*⟩ ^2^, ⟨*Y* ⟩ ⟨*Z*⟩, and max(⟨*Y* ⟩, ⟨*Z*⟩ ), which probe sub-linear, super-linear, multiplicative, and bottleneck regimes; and the single-species costs ⟨*Y* ⟩ and ⟨*Z*⟩ . Relative to the baseline, the within-family ranking of variants by Pareto share has mean Spearman correlation 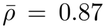 across the linear weights, with the lowest value, 0.61, at *α/β* = 1*/*3, where the output dominates the cost, and 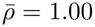 for all four nonlinear and both single-species costs. The winning set {C1, C3, I1, I3} is preserved in all seven linear weightings, including the extreme *α/β* = 1*/*3, so the excluded variants remain dominated under every cost specification tested.

**TABLE I.**
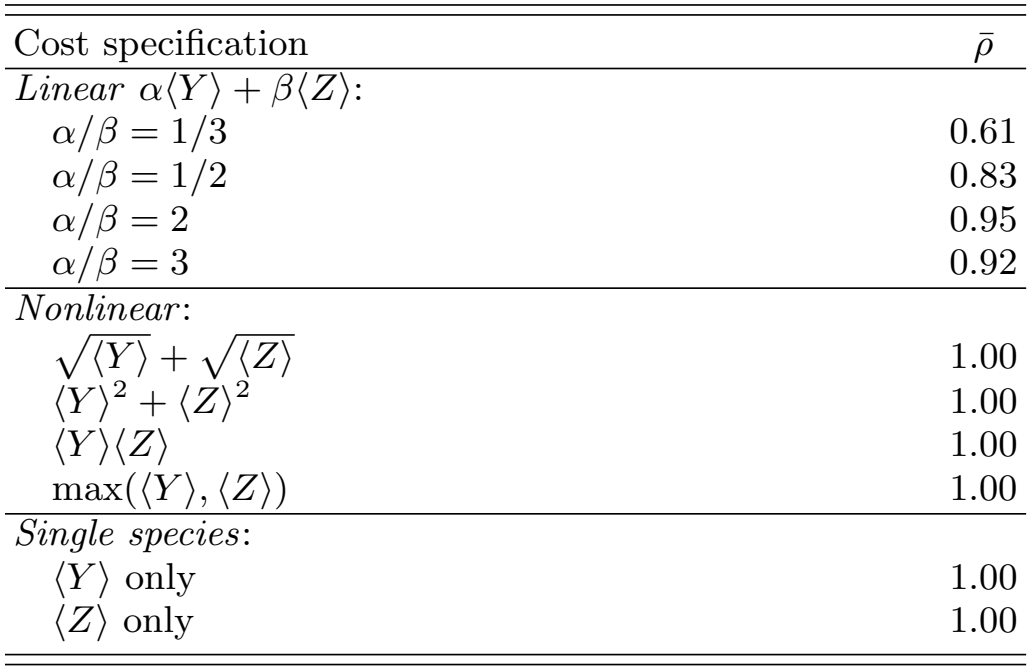
Robustness of the within-family Pareto-share ranking to the cost specification. Each row is the mean Spearman correlation of the ranking with the baseline *B* = ⟨*Y* ⟩ + ⟨*Z*⟩, averaged over the eight (family, gate, *n*) subgroups, on the *N* = 1000 samples. The ranking is unchanged 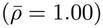 under every nonlinear and single-species cost, and departs from the baseline only for linear weights that heavily favor the output.

| Cost specification | ρ̄ |
|---|---|
| *Linear* α⟨Y⟩ + β⟨Z⟩: |  |
| α/β = 1/3 | 0.61 |
| α/β = 1/2 | 0.83 |
| α/β = 2 | 0.95 |
| α/β = 3 | 0.92 |
| *Nonlinear*: |  |
| √⟨Y⟩ + √⟨Z⟩ | 1.00 |
| ⟨Y⟩<sup>2</sup> + ⟨Z⟩<sup>2</sup> | 1.00 |
| ⟨Y⟩⟨Z⟩ | 1.00 |
| max(⟨Y⟩, ⟨Z⟩) | 1.00 |
| *Single species*: |  |
| ⟨Y⟩ only | 1.00 |
| ⟨Z⟩ only | 1.00 |

**TABLE II.** Within-family rankings by mean information over *N* = 1000 parameter samples, from the analytical capacity *C* and from the Gillespie estimate *Î*_SSA_(*E*; *Z*), with their Spearman correlation. The rankings coincide in five subgroups and differ by one adjacent transposition in the other three, all incoherent. These are rankings by information alone; the Pareto shares of Sec. VI B also account for the burden *B* and order the variants differently.

| Family | Gate | *n* | Spearman *ρ* | Lyapunov ranking | SSA ranking |
|---|---|---|---|---|---|
| Coherent | OR | 2 | 1.00 | C1 > C2 > C4 > C3 | C1 > C2 > C4 > C3 |
| Coherent | AND | 2 | 1.00 | C3 > C2 > C4 > C1 | C3 > C2 > C4 > C1 |
| Coherent | OR | 4 | 1.00 | C1 > C2 > C4 > C3 | C1 > C2 > C4 > C3 |
| Coherent | AND | 4 | 1.00 | C3 > C2 > C4 > C1 | C3 > C2 > C4 > C1 |
| Incoherent | OR | 2 | 0.80 | I3 > I2 > I1 > I4 | I3 > I2 > I4 > I1 |
| Incoherent | AND | 2 | 1.00 | I1 > I2 > I3 > I4 | I1 > I2 > I3 > I4 |
| Incoherent | OR | 4 | 0.80 | I2 > I3 > I4 > I1 | I3 > I2 > I4 > I1 |
| Incoherent | AND | 4 | 0.80 | I1 > I2 > I4 > I3 | I1 > I4 > I2 > I3 |

**TABLE III.** Stratified Pareto shares (%) with 95% bootstrap intervals over 2000 resamples of the *N* = 1000 parameter samples, and Wilcoxon signed-rank tests on the per-sample difference *D*_*k*_ = share(I1)_*k*_ − share(I3)_*k*_, with the matched-pairs rank-biserial effect size *r*_rb_ (the rank correlation of the signed differences, bounded in [−1, 1]) and the number of non-tied pairs. The *p*–values are driven by the Monte Carlo sample size *N* = 1000 and quantify resolution rather than biological effect size; the argument rests on the effect sizes *r*_rb_ and the bootstrap intervals, not on the *p*–values. Shares are stratified within family, so the four coherent shares and the four incoherent shares each sum to 100%. C1 leads the coherent family in all four subgroups, with its lower interval bound above the upper bound of every other coherent variant. I3 lies above I1 in the three subgroups where the paired test survives Holm-Bonferroni correction; the two overlap only for (AND, *n* = 4). The robustness of these stationary differences to low-information samples is examined in Sec. VI C.

| Subgroup | C1 (%) | C2 (%) | C3 (%) | C4 (%) |
|---|---|---|---|---|
| OR, n=2 | 51.7 [48.8, 54.5] | 17.6 [15.5, 19.8] | 9.2 [7.6, 10.8] | 21.5 [19.3, 23.9] |
| AND, n=2 | 32.2 [30.8, 33.5] | 23.7 [22.2, 25.2] | 23.9 [22.4, 25.5] | 20.2 [18.7, 21.8] |
| OR, n=4 | 49.6 [47.1, 52.3] | 19.4 [17.5, 21.6] | 7.8 [6.5, 9.2] | 23.1 [20.7, 25.4] |
| AND, n=4 | 30.1 [28.9, 31.3] | 22.7 [21.4, 24.1] | 25.3 [24.0, 26.7] | 21.9 [20.6, 23.1] |
| Subgroup | I1 (%) | I2 (%) | I3 (%) | I4 (%) |
| OR, n=2 | 21.3 [19.1, 23.6] | 28.2 [25.7, 30.8] | 35.0 [32.4, 37.8] | 15.5 [13.7, 17.2] |
| AND, n=2 | 26.9 [25.6, 28.2] | 25.1 [23.9, 26.4] | 30.4 [29.1, 31.7] | 17.6 [16.2, 18.9] |
| OR, n=4 | 17.0 [15.1, 18.9] | 33.2 [30.6, 35.9] | 37.4 [34.7, 40.1] | 12.5 [10.8, 14.1] |
| AND, n=4 | 28.4 [27.1, 29.7] | 22.4 [21.2, 23.5] | 30.2 [29.1, 31.4] | 19.0 [17.8, 20.3] |
| **Paired Wilcoxon test on** *D*<sub>*k*</sub> = share(I1)<sub>*k*</sub> − share(I3)<sub>*k*</sub> **(Holm-corrected):** |  |  |  |  |
| OR, n=2 | *p*<sub>Holm</sub> = 1.6 × 10<sup>−10</sup>; mean *D*<sub>*k*</sub> = −0.137; *r*<sub>rb</sub> = −0.27; 730 pairs |  |  |  |
| AND, n=2 | *p*<sub>Holm</sub> = 9.2 × 10<sup>−5</sup>; mean *D*<sub>*k*</sub> = −0.036; *r*<sub>rb</sub> = −0.26; 300 pairs |  |  |  |
| OR, n=4 | *p*<sub>Holm</sub> = 2.6 × 10<sup>−22</sup>; mean *D*<sub>*k*</sub> = −0.204; *r*<sub>rb</sub> = −0.42; 699 pairs |  |  |  |
| AND, n=4 | *p*<sub>Holm</sub> = 0.088; mean *D*<sub>*k*</sub> = −0.018; *r*<sub>rb</sub> = −0.11; 275 pairs |  |  |  |

**TABLE IV.** Per-subgroup linear fit *C* = *a Î*_SSA_ + *b* (clipped *C* = 0 samples excluded) and Pearson correlation, with 95% bootstrap intervals on the slope over 2000 resamples of the *N* = 1000 paired measurements, and the fraction of samples clipped to *C* = 0 in each subgroup. The slopes do not separate the two families (Mann-Whitney *p* = 0.49), so the offset carries no systematic branch-degeneracy signature. Clipping is concentrated in the incoherent OR subgroups, where the non-monotonic response most often leaves the small-noise integral below its floor.

| Family | Gate | *n* | Slope *a* | Interc. *b* | Pearson *r* | clip |
|---|---|---|---|---|---|---|
| Coherent | OR | 2 | 1.23 [1.21, 1.26] | −0.26 | 0.95 | 20.5% |
| Coherent | AND | 2 | 1.09 [1.06, 1.12] | +0.01 | 0.93 | 2.8% |
| Coherent | OR | 4 | 0.92 [0.89, 0.94] | +0.27 | 0.88 | 27.5% |
| Coherent | AND | 4 | 0.82 [0.79, 0.84] | +0.55 | 0.83 | 13.2% |
| Incoherent | OR | 2 | 1.22 [1.18, 1.25] | −0.15 | 0.83 | 44.5% |
| Incoherent | AND | 2 | 1.10 [1.07, 1.13] | +0.12 | 0.82 | 15.2% |
| Incoherent | OR | 4 | 1.16 [1.13, 1.20] | −0.01 | 0.88 | 47.7% |
| Incoherent | AND | 4 | 0.98 [0.95, 1.02] | +0.43 | 0.80 | 23.2% |

Because dominance depends only on the order of costs, the frontier is invariant under any strictly increasing transformation *B* ↦*ϕ*(*B*). The nonlinear costs above are not such transformations but distinct functions of (⟨*Y* ⟩, ⟨*Z*⟩ ), so their agreement with the baseline is an empirical result rather than a consequence of this invariance. Costs that include mRNA, *B*^′^ = *B* + *λ*(⟨*m*_Y_⟩ +⟨ *m*_Z_ ⟩), reduce at deterministic steady state to a linear weighting of ⟨*Y* ⟩ and ⟨*Z*⟩, since ⟨*m*_i_⟩ = (*µ*_i_*/K*_pi_) ⟨*i*⟩ ; with the node-independent rates of App. B, *B*^′^ is proportional to *B* and leaves the frontier unchanged.

### D. Robustness to the kinetic-parameter prior

The Pareto shares depend on the log-uniform sampling distribution of (*β*_y_, *β*_z_, *K*_XY_, *K*_XZ_, *K*_Y Z_) (App. B). This reference distribution weights all configurations equally in log-space, whereas real parameter distributions are shaped by regulatory context and evolutionary history [32]. We therefore reweighted the *N* = 1000 existing samples, without new simulations, with two alternative priors: a Gaussian prior in log-space centered on typical values from proteome-wide measurements [32] and dissociation-constant ranges in BioNumbers [33], and an exponential weighting that favors low-expression samples, as in strongly regulated stress and sensory genes. Under both priors the within-family ranking keeps Spearman *ρ >* 0.85 relative to the baseline in all (gate, *n*) sub-groups, while individual shares move by up to about 10– 15 percentage points. Reweighting a fixed sample is reliable only if the new prior does not concentrate where the original draws are sparse. The Kish effective sample sizes, 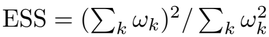 with *ω*_k_ the sample weights, are 337 (67% of *N* ) for the Gaussian prior and 344 (69%) for the exponential prior, above the ESS*/N*≈ 0.5 level below which reweighting becomes unreliable. The rankings are therefore not an artifact of the reference distribution, although neither alternative is a measured parameter distribution.

## VI. RESULTS

We now apply the framework to the eight FFL variants. Degradation and translation rates are equal across nodes and fixed (App. B), and time is measured in units of the mRNA lifetime (*γ* = 1). The parameters (*β*_y_, *β*_z_, *K*_XY_, *K*_XZ_, *K*_Y Z_) are sampled log-uniformly, and *n* ∈ {2, 4}.

### A. Local mutual information and channel capacity

Figure 2 shows, for each variant at *n* = 2 under OR gating, the analytical dose-response ⟨*Z*⟩(*E*) from the Lyapunov solution of Eq. (18) with its conditional noise *σ*_Z_(*E*), and, on the same panel, the local Gaussian mutual information *I*_loc_(*E*) of Eq. (21). The four coherent variants occupy the top row and the four incoherent variants the bottom row, on a common *E* axis.

**FIG 2.**
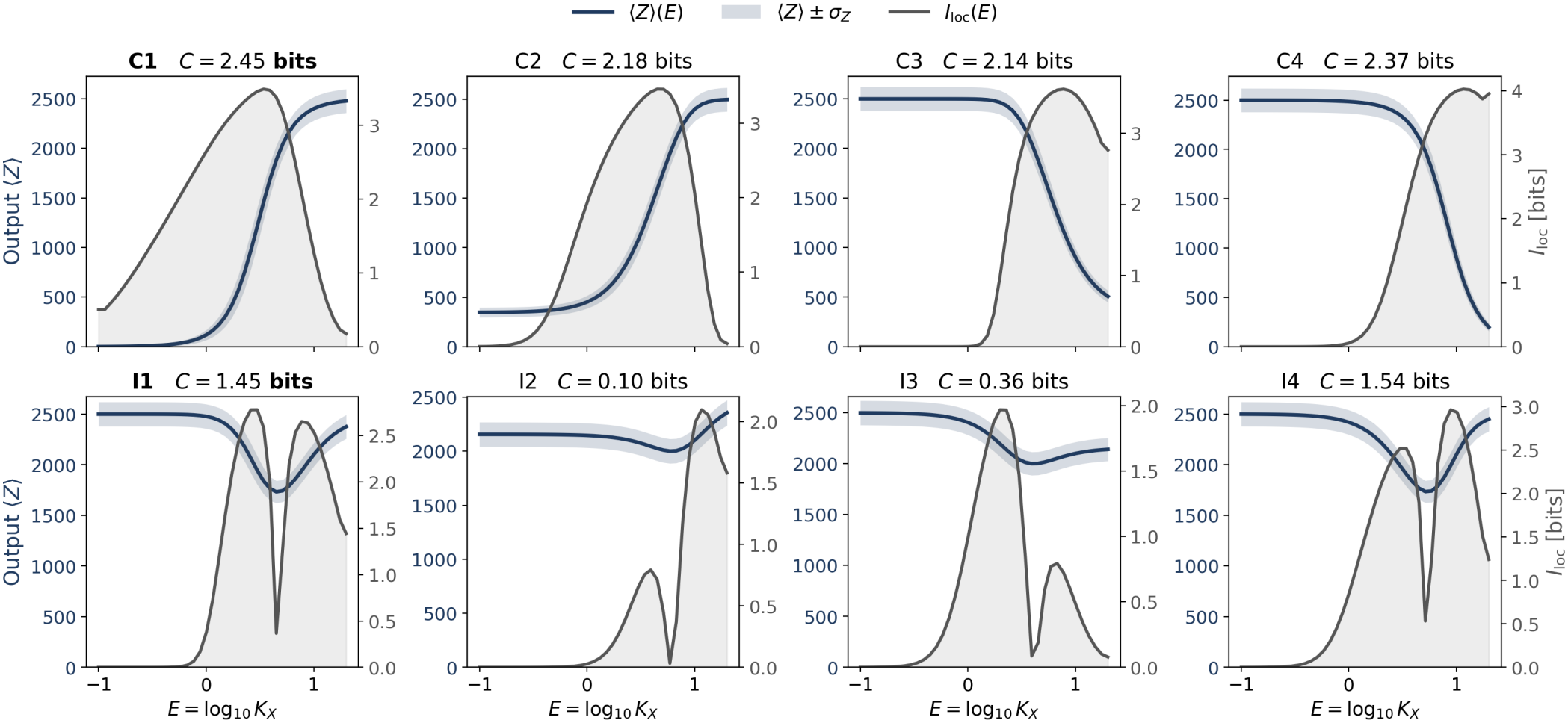
Analytical dose-response and local mutual information for the eight FFL variants under OR gating at Hill coefficient *n* = 2, computed at a representative parameter point. Each panel shows, on a twin axis, the mean output ⟨*Z*⟩(*E*) (navy line, left axis) with the conditional noise as a shaded band ⟨*Z*⟩ ± *σ*_*Z*_(*E*), and the local Gaussian mutual information *I*_loc_(*E*) of Eq. (21) (grey line, right axis), as a function of *E* = log_10_ *K*_*X*_ . The integrated channel capacity *C* of Eq. (22) is printed in each panel title. The coherent variants (C1–C4, top row) have monotonic dose-responses and single-peaked *I*_loc_ profiles; the incoherent variants (I1–I4, bottom row) have non-monotonic, band-pass dose-responses and characteristic two-peaked *I*_loc_ profiles separated by the valley where the gain vanishes, the analytical signature of band-pass information transmission.

Two features stand out. The coherent variants have monotonic dose-responses, increasing for C1 and C2 and decreasing for C3 and C4, and therefore single-peaked *I*_loc_(*E*) profiles centered on the sigmoidal transition, where the response is steepest and the circuit best distinguishes neighboring environments. The incoherent variants have non-monotonic, band-pass dose-responses with a maximum at intermediate *E*, so *I*_loc_(*E*) has two peaks separated by a valley at the maximum, where the gain *g*(*E*) vanishes and no local information is transmitted. This qualitative difference in the shape of the information profile, one peak against two, is the first indication that the two families realize distinct functions and motivates comparing architectures within a family rather than across families (Sec. V). The two-peaked profile is the steady-state signature of the band-pass behavior of inco-herent FFLs, whose dynamical counterpart is the pulse generation described in deterministic analyses of I1 [24]; it is also the origin of the branch degeneracy addressed in Sec. IV D, since the two environmental states on either side of the maximum share the same mean output.

The capacity *C* summarizes each curve in a single number. At this point C1 has the highest coherent capacity (2.45 bits, against 2.14–2.37 bits for the other coherents), I1 and I4 reach 1.45 and 1.54 bits, and I2 and I3 only 0.10 and 0.36 bits, because their gain stays small across the operational range. These values describe a single parameter point and do not rank the variants in general. Averaged over parameter samples (Table II), the ordering by information alone depends on the gate: under OR, C1 leads the coherents and I3 or I2 the incoherents, whereas under AND, C1 has the lowest mean information among the coherents and I1 leads the incoherents. Information rankings are also not cost-information rankings. A variant with little information can lie on the Pareto frontier if its burden is correspondingly low: at the point of Fig. 2, I3 has a much lower burden *B* than I1, so the two variants are mutually non-dominated despite their fourfold capacity gap. The cost axis likewise allows C1 to lead the coherent Pareto shares under both gates (Sec. VI B) although it does not lead in mean information under AND.

### B. Stationary Pareto shares from stochastic simulations

We next examine the dependence on parameters. Figure 3 shows the stratified Pareto shares of the coherent and incoherent variants at *n* = 2 and 4 under OR and AND gating, computed from *N* = 1000 parameter samples with the stationary mutual information estimated from Gillespie simulations.

**FIG 3.**
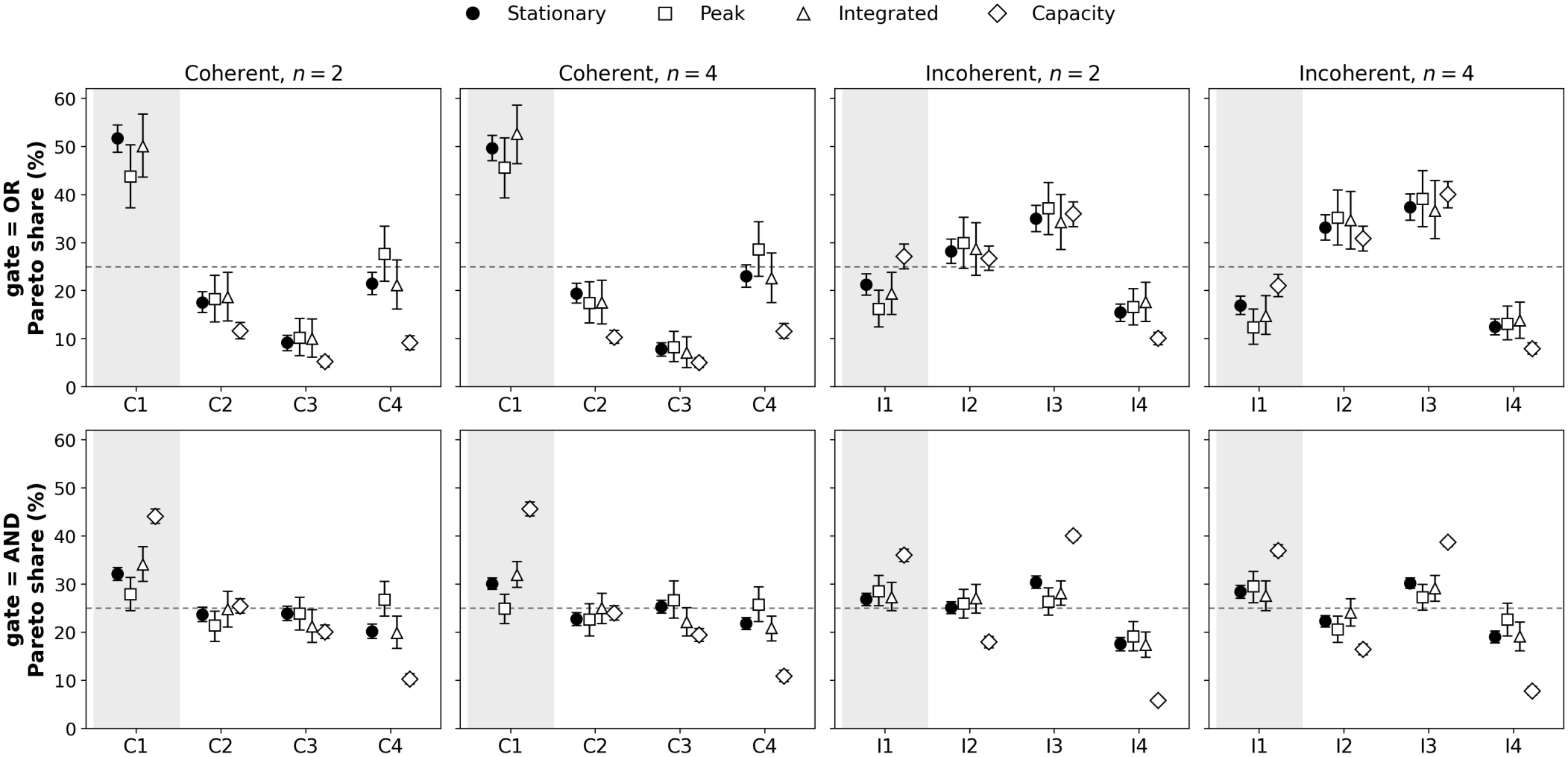
Stratified Pareto shares of the four within-family variants under the four information measures: stationary (filled circle), peak transient (open square), and time-integrated (open triangle) mutual information, and analytical channel capacity (open diamond). Rows are the OR and AND gates; columns are the two families at *n* = 2 and 4. At each parameter sample, every variant on the cost-information frontier receives credit 1*/k* for a frontier of *k* variants, and credits are averaged over samples (*N* = 1000 for the stationary and capacity measures, *N* = 200 for the two transient measures). Error bars are 95% bootstrap intervals; the shaded column marks the empirically dominant variant (C1, I1) and the dashed line the uniform 25% level. Under OR the four measures agree that C1 leads the coherent family and that I3 exceeds I1; under AND the shares are closer to uniform and I1 and I3 are comparable. Because frontiers typically contain several variants, all members of a family carry appreciable share; the four-variant reduction appears in the sole-winner analysis (Fig. 4) and in the unstratified comparison (Fig. 5). Bootstrap intervals for the stationary measure are listed in Table III.

Three observations follow. First, the shares are fairly flat within each family: every variant carries between about 8% and 51% and none is at zero, because the frontier at a typical sample contains several variants. The stratified Pareto share alone therefore does not single out a four-variant set; that reduction comes from the sole-winner analysis of Sec. VI F and the unstratified comparison of Sec. VI G. Second, C1 is the only coherent variant consistently above the uniform 25% level, with shares from 30.1% (AND, *n* = 4) to 51.7% (OR, *n* = 2), and its 95% bootstrap interval lies above every other coherent variant in all four subgroups (Table III). The nearest competitor depends on the gate: under OR it is C4, and under AND the share is spread almost evenly over C2, C3, and C4 near the uniform 25% level (Fig. 3). Across all ten gates (App. C 4) C1 leads under six logics but is displaced by C3 under the NAND–, NOR–, XOR– and XNOR-like logics. Third, no incoherent variant plays the role of C1: I3 (30.2%–37.4%), I2 (22.4%–33.2%), and I1 (17.0%–28.4%) all carry substantial share.

To test whether I1 and I3 are resolved, we computed the per-sample difference *D*_k_ = share(I1)_k_ − share(I3)_k_ and applied a Wilcoxon signed-rank test in each of the four (gate, *n*) subgroups, with Holm-Bonferroni correction for the four tests (Table III). Under OR at both *n* and under AND at *n* = 2, the test favors I3 with *p*_Holm_ *<* 10^−4^: the mean *D*_k_ ranges from − 0.036 (AND, *n* = 2) to − 0.204 (OR, *n* = 4), with rank-biserial effect sizes from −0.26 to −0.42, and I3 carries the larger share in 438 of the 730 non-tied samples under (OR, *n* = 2) and in 457 of 699 under (OR, *n* = 4). Under (AND, *n* = 4) the difference is not significant (*p*_Holm_ = 0.09). Many samples place I1 and I3 exactly level, so the median *D*_k_ is zero throughout.

The direction of this stationary difference is reproduced by the other three measures under OR but not under AND, and it is robust to removing non-responsive samples (Sec. VI C). The peak transient, time-integrated, and analytical capacity measures, the last with *N* = 1000 samples, reproduce the I3 lead under OR and return an I1/I3 tie under AND (Secs. VI D and VI E). Across these share measures the lead, where there is one, is I3’s; the sole-winner mass leans the other way by a comparable margin (Sec. VI F), so the two variants are close to equivalent under the trade-off. The implications for the census, which favors I1 far more strongly, are discussed in Sec. VII.

### C. Robustness to low-information samples

The log-uniform prior of App. B weights kinetic parameters over wide absolute ranges and therefore includes combinations in which a variant barely responds to the environment over the operational range of *E*, so that it transmits little information at any cost (the choice of prior is discussed in App. B). The concern is that a variant sitting near zero information is placed on the frontier by its burden alone, so that a cheap, uninformative architecture could accumulate Pareto share for the wrong reason. We address this with two filters that both preserve the paired four-variant comparison, using the SSA estimates of Sec. IV D and sweeping an information threshold from 0 to 0.5 bits for all ten gates and both Hill coefficients.

The first, weaker filter discards a family only when *no* variant in it reaches the threshold, removing the samples where all four are dead. Under this filter the rankings are unchanged across the whole threshold range: the coherent leader of each gate (C1 under most logics, C3 under the NAND–, NOR–, XOR– and XNOR-like ones) is preserved, C1 holds its canonical-gate lead essentially unchanged up to 0.5 bits, and the mean *D*_k_ = share(I1) − share(I3) keeps its value (for example, from −0.137 to −0.139 under OR at *n* = 2). This filter, however, only removes wholly dead samples, which are rare; a typical retained sample may still contain two or three near-zero variants competing on cost, so it does not fully address the concern.

The second, stricter filter is the direct test: it retains a family only when *every* variant reaches the threshold, so the surviving samples contain four informative variants competing on a fixed four-element frontier, with the burden-only ties removed and the pairing intact (Fig. 8 in App. C 5). The outcome is gate-dependent, for a mechanical reason: under OR the incoherent variants I2 and I3 are the ones most often in the unresponsive regime, so requiring all four to be informative is rarely satisfied and the retained sample collapses (from 1000 to 85–130 at 0.1–0.2 bits), leaving too few samples to resolve *D*_k_; the bootstrap interval on *D*_k_ then widens to include zero, which reflects loss of power rather than equivalence. Under AND the four variants respond more evenly and the filter retains 640 to 900 samples. There the result is decisive and shows no reversal: at *n* = 2 the mean *D*_k_ stays at −0.035 to −0.039 with a bootstrap interval that excludes zero at every threshold, and at *n* = 4 the already marginal difference (mean *D*_k_ = −0.018, *p*_Holm_ = 0.088 without filtering) relaxes toward zero with an interval that includes it, never crossing to favor I1. The concern that I3 accumulates share under AND by being cheap where it transmits nothing is therefore not borne out: removing the uninformative samples, with the pairing preserved, does not make I1 emerge.

We note for completeness that a variant-level filter, which drops individual variants below the threshold rather than whole families, does produce an apparent I1–favoring reversal under AND at *n* = 4. That filter changes the size of the comparison set and hence the 1*/k* credit and breaks the pairing, so its reversal is an accounting artifact rather than a property of the trade-off; the common-support filter above, which keeps the four-element set fixed, supersedes it. Across both valid filters no measure separates I1 from I3 by more than a few percentage points, consistent with the near-equivalence of the two variants.

### D. Dynamic Pareto shares

The stationary measure ignores timing, which is central to the function of FFLs: in deterministic analyses, coherent and incoherent variants differ mainly in the shape of their transient response [24]. We therefore performed step-response experiments. Each system is equilibrated at 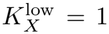 and, at *t* = 0, switched to 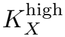, one of eight values log-spaced in [1, 8]. These eight separate step experiments define a uniform prior over 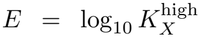 . From the eight conditional distributions 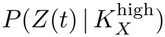 we compute *I*(*E*; *Z*(*t*)) at each post-step time and extract the peak information *I*_peak_ = max_t_ *I*(*E*; *Z*(*t*)) and the time-integrated information 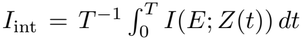. Both still quantify information about the *level* of the new environment, read out at or averaged over post-step times; neither measures information about *when* the switch occurred, so all four of our measures are level measures, not timing measures.

The corresponding burdens are *B*_peak_, evaluated at the time of the peak, and *B*_int_, averaged over [0, *T* ]. The cost of these simulations limits the scan to *N* = 200 parameter samples with *R* = 20 replicates each, so the dynamic shares carry a larger sampling uncertainty than the stationary ones and we interpret them qualitatively.

The resulting Pareto shares (Fig. 3, open squares and triangles) reproduce the coherent structure of the stationary analysis, with C1 leading. They also reproduce the gate dependence of the I1/I3 relation: under both peak and time-integrated information, I3 leads I1 under OR (mean *D*_k_ from −0.15 to −0.27, *p <* 10^−3^) while the two are indistinguishable under AND. No dynamic measure favors I1 over I3.

### E. Analytical Pareto shares from the channel capacity

The Gillespie analyses are computationally expensive and use a discrete uniform prior. The capacity of Eq. (22), by contrast, takes a fraction of a second per parameter sample and integrates over the whole operational range under the optimal prior. We used this speed to compute Pareto shares from *C* and *B* at *N* = 1000 samples (Fig. 3, open diamonds). C1 again leads the coherent family, with a larger margin (73% of the share under OR and 44%–46% under AND). Within the incoherent family, I3 leads I1 under OR (mean *D*_k_ = − 0.19 at *n* = 4, *p <* 10^−14^) and the two are close under AND, matching the stationary and dynamic measures. The coherent ranking thus agrees between an empirical estimator with a discrete uniform prior and an analytical capacity with an optimal continuous prior. In the Pareto shares, I3 lies at or above I1 under every measure; the sole-winner mass instead gives I1 a narrow edge (Sec. VI F), and neither separation exceeds a few percentage points.

### F.Robustness across gate logics: the gate simplex

The analyses above use the AND and OR gates. We now include all ten gates through the mixture analysis of Sec. V B. Figure 4 shows the sole-winner mass of each variant for the four subgroups (family × *n*) and the four information measures. C1 is the most frequent coherent winner in every subgroup and under every measure (46%– 80% of the simplex), and C3 is the only other coherent variant that ever wins. I1 and I3 share all of the inco-herent sole-winner mass, in proportions that vary across measures, and I2 and I4 never win. The four variants C1, C3, I1, and I3 therefore absorb all of the sole-winner mass, and C2, C4, I2, and I4 have exactly zero under every measure. The measures differ in the exact percentages but agree on this four-variant set.

**FIG 4.**
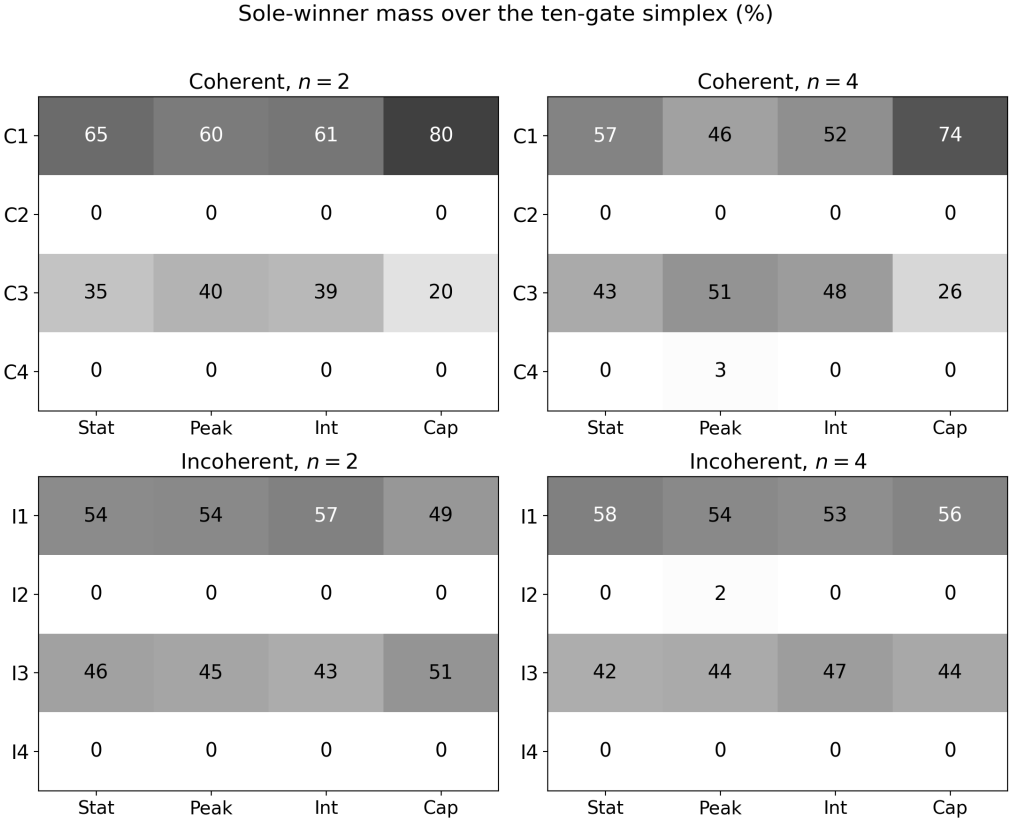
Sole-winner mass (%) over the ten-gate simplex, shown as a greyscale map of variant (rows) against information measure (columns): stationary, peak transient, and time-integrated SSA mutual information, and analytical capacity. Each cell is the fraction of the Dirichlet simplex Δ^9^ on which that variant has the largest gate-mixed Pareto share within its family; the four panels are the family × *n* subgroups. C1 is the most frequent coherent winner in every panel and C3 the only other coherent winner; I1 and I3 share all of the incoherent mass. The rows for C2, C4, I2, and I4 are identically zero, which is the four-variant reduction.

The incoherent sole-winner mass is close to evenly divided between I1 and I3 (I1 takes 49%–58% of the incoherent mass across the eight subgroup and measure combinations, a near-even split), and its slight lead for I1 runs opposite to the stratified Pareto shares, which lead for I3 (Sec. VI B). The two statistics measure different things and need not agree: the Pareto share rewards a variant for lying on the frontier across many gates, so it favors the variant that is consistently competitive, whereas the sole-winner mass rewards a variant for the size of its lead where it does win, so it favors the variant whose advantage is concentrated in a few gates with large margins. I3 is competitive under most gates without a decisive lead in any, while I1 wins fewer gates but by wider margins; the first pattern accumulates Pareto share and the second accumulates sole-winner mass. Neither statistic separates I1 from I3 by more than a few percentage points, and they disagree on the sign of that small difference, so the incoherent comparison is close to a tie under both.

### G. Unstratified eight-variant comparison

As a control on stratification, all eight variants compete in a single Pareto comparison at each sample (Fig. 5); the shares then sum to unity over eight variants, and the uniform level is 12.5%. Only C1, C3, I1, and I3 carry non-negligible share, while C2, C4, I2, and I4 are at or near zero in all four (gate, *n*) combinations. This is the four-variant set of the sole-winner analysis, here obtained without stratifying by family. C1 carries the largest share (28%–56%), followed by I1 or I3 depending on the gate, with C3 typically fourth. Removing the stratification therefore changes which variants carry frontier mass: C2, C4, I2, and I4 hold appreciable shares within their own family (Fig. 3) but are dominated once both families compete. We use the stratified analysis in the main text for the biological reasons given in Sec. V.

**FIG 5.**
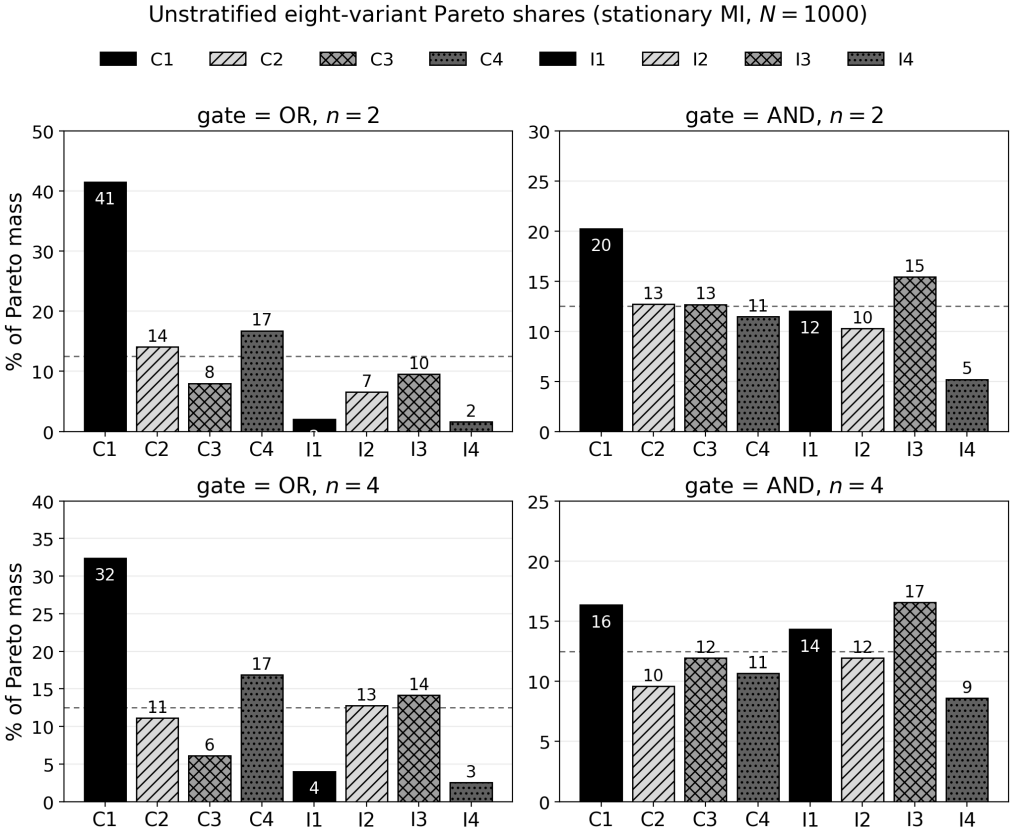
Unstratified eight-variant Pareto shares under OR and AND gating at *n* = 2 and 4: all eight variants compete on a single frontier and the shares sum to 100%, with the dashed line at the uniform 12.5% level. Bars are filled in greyscale with a distinct hatch per variant and a solid black bar for the dominant variant (C1, I1). Even when coherent and incoherent variants compete directly, C1, C3, I1, and I3 carry essentially all of the share, and C2, C4, I2, and I4 fall to or near zero.

## VII. DISCUSSION

We have compared the eight FFL variants along two axes, the environment-to-output information they transmit and the protein burden *B* they pay, using an analytical capacity derived from the linearized chemical Langevin dynamics and mutual information estimated from Gillespie simulations at steady state and after step inputs. The two estimators agree closely (Sec. IV D), and the within-family rankings are largely insensitive to the cost specification and to the parameter prior (Secs. V C and V D).

The status of the comparison with the census should be stated explicitly. Partial agreement between what the framework selects and what is abundant in *E. coli* and *S. cerevisiae* indicates that the cost-information trade-off is a plausible contribution to the census, not that it is the only or the dominant selection pressure. The map from any performance metric to motif abundance also passes through the fitness landscape, mutational accessibility, the embedding of each FFL in larger modules, correlations among upstream regulators, and the environmental statistics of each organism, none of which our analysis constrains. Appendix C 6 (Table V) lists the empirical counts of Ref. [21] alongside the stationary Pareto shares. The shares are nonzero for all eight variants and so do not by themselves carry the four-variant reduction, which follows instead from the sole-winner and unstrati-fied analyses.

**TABLE V.** Empirical FFL counts from Ref. [21] for *E. coli* (42 FFLs) and *S. cerevisiae* (56 FFLs), compared with the stratified Pareto shares of the stationary analysis at *n* = 2 and OR gating, with 95% bootstrap intervals over 2000 resamples of the *N* = 1000 parameter samples. The shares measure how often a variant lies on the cost-information frontier and are nonzero for all eight variants; the four-variant reduction is a property of the sole-winner analysis (Fig. 4), not of these shares.

| Motif | *E. coli*<br>count | *S. cer.*<br>count | Pareto share<br>(%) [95% CI] |
|---|---|---|---|
| C1 | 28 | 26 | 51.7 [48.8, 54.5] |
| C2 | 2 | 5 | 17.6 [15.5, 19.8] |
| C3 | 4 | 0 | 9.2 [7.6, 10.8] |
| C4 | 1 | 0 | 21.5 [19.3, 23.9] |
| I1 | 5 | 21 | 21.3 [19.1, 23.6] |
| I2 | 0 | 3 | 28.2 [25.7, 30.8] |
| I3 | 1 | 1 | 35.0 [32.4, 37.8] |
| I4 | 1 | 0 | 15.5 [13.7, 17.2] |

Three findings emerge. First, only C1, C3, I1, and I3 are ever the sole within-family winner at any point of the ten-gate simplex, under all four information measures, and C2, C4, I2, and I4 have exactly zero sole-winner mass (Fig. 4); the unstratified comparison recovers the same set (Sec. VI G). The excluded variants are rare in both organisms [21]. The match is partial, however, because C3 and I3 are rare too: C3 appears 4 times in *E. coli* and never in *S. cerevisiae*, and I3 once in each, whereas in yeast the excluded C2 (5) and I2 (3) are more frequent than either. The framework thus selects C3 and I3 in addition to the abundant C1 and I1. At counts of 0, 1, and 4, however, the census cannot distinguish a genuine exclusion of C3 and I3 from sampling noise, so we do not read them as clear failures: the resolved structural claim is the exclusion of the four variants that never win, C2, C4, I2, and I4, which are rare in both organisms. Whether the selection of C3 and I3 reflects a real over-selection or the limited resolution of the census cannot be settled with these numbers.

Second, C1 leads the coherent family in all four (gate, *n*) combinations, with mean stationary shares between 30% and 51% and 95% bootstrap intervals above every other coherent variant (Table III), and it also leads under the peak transient, time-integrated, and analytical capacity measures. Its lead over the coherent runner-up shrinks from about 30 percentage points under (OR, *n* = 2), where the other three variants sit far below, to a few percentage points under (AND, *n* = 4), where the coherent shares are nearly uniform, so the dominance is uniform in sign but graded in magnitude. This agrees with the census, in which C1 is the most abundant coherent variant in both organisms [21].

Third, no information measure separates I1 from I3 by anything close to the margin of the census. The stratified Pareto shares and the two transient measures place I3 slightly ahead of I1 under OR (Holm-corrected *p <* 10^−3^; mean per-sample share differences from 0.14 to − 0.27), and the analytical capacity does the same; under AND the two are statistically indistinguishable. The sole-winner mass over the gate simplex leans the other way, giving I1 a narrow share (49%–58% of the incoherent mass across measures, a near-even split), because the two statistics weight the frontier differently (Sec. VI F). By individual gate the lead is split evenly, I1 ahead under five of the ten logics and I3 under the other five (App. C 4). No statistic resolves the incoherent pair by more than a few percentage points, and they disagree on the sign of that difference, so the framework treats I1 and I3 as nearly equivalent. In the census, by contrast, I1 outnumbers I3 by 21:1 in *S. cerevisiae*; the 5:1 ratio in *E. coli* is consistent with chance at these counts, so the significant asymmetry is the one in yeast. Even that asymmetry cannot be attributed to cost and information, which are close to indifferent between the two. Our four measures all quantify information about the level of the environment, however, not about the timing of its changes, and the functional literature attributes the advantage of I1 to response acceleration and pulse generation, both properties of timing. The separating axis may therefore lie in a timing-sensitive measure, which our level-information measures do not probe, rather than strictly outside the cost-information plane; distinguishing these requires the extension noted below. We do not claim that selection opposes a cost-information gradient.

This division between the two families is consistent with the functional literature. The proposed functions of both dominant variants are dynamical: persistence detection by C1 [22, 23], and response acceleration and pulse generation by I1 [24, 25]. The difference revealed here is that C1 is also favored by cost and information, whereas an advantage of I1 would have to rest on properties our measures do not resolve, such as the shape and timing of the transient, or on other axes. The incoherent architecture produces the band-pass dose-response and two-peaked *I*_loc_(*E*) of Fig. 2 and places I1 on the dynamic frontiers, but it does the same for I3. That cost and information fail to favor I1 is thus consistent with the emphasis of the literature on the dynamics of I1, and with environmental-selection arguments that depend on the statistics of the input [26, 27]. What our analysis adds is a parameter-averaged ranking of all eight variants on a common information and burden axis; what it cannot identify is the axis along which selection separates I1 from I3.

Several modeling choices limit these conclusions. First, the Pareto shares are defined relative to a log-uniform reference distribution of kinetic parameters. Reweighting to two alternative priors preserves the rankings (Sec. V D), but neither alternative is a measured distribution, so the shares are not predictions of motif fractions in any organism. Second, the analytical capacity relies on the linear-noise and small-noise approximations. The Gillespie estimates do not, and the two agree over our parameter range (Sec. IV D), but the approximations may fail at very low copy numbers or under strong non-linearity [12, 36], and fundamental limits on noise suppression [15] bound the cost of fidelity for any architecture. Third, the protein-burden proxy is a coarse representation of the cost of expression; the rankings survive the alternatives of Sec. V C, but the percentages shift. Fourth, degradation and translation rates are equal across nodes and fixed, as is the timescale separation between mRNA and protein (App. B). This isolates the effect of topology, but node-asymmetric degradation could change the balance between ⟨*Y* ⟩ and ⟨*Z*⟩ in the burden. Given the cost-robustness results, we expect the within-family ordering to tolerate moderate asymmetry, but we have not tested strongly asymmetric kinetics. Fifth, the model includes only intrinsic reaction noise: extrinsic noise, cell-to-cell variability in the environmental input *E*, and any time-averaging by downstream readouts are absent [53], and all three are known to compress the capacity differences between architectures, so the differences reported here are, if anything, upper bounds on what a noisier readout would resolve. Finally, the comparison is restricted to the two organisms with well-characterized censuses, and we do not claim that it transfers unchanged to networks with substantially different noise and timescale regimes, such as those of multicellular eukaryotes.

The analysis belongs to a broader programme in which information theory and non-equilibrium statistical mechanics are used to rationalize the architecture of biological circuits [8, 10–14]. Its results suggest that part of the apparent diversity of network motifs can be organized by a small number of physical trade-offs, and also that such trade-offs leave part of the census unexplained. Three extensions could test this picture: applying the framework to four-node motifs such as bi-fans, single-input modules, and dense overlapping regulons [2, 19]; extending the information measures to the timing of environmental changes, for instance the mutual information between the switch time and *Z*(*t*) at a fixed observation time, or a burden that charges for response delay, which is a direct extension of the step-response machinery of Sec. VI D and would test the timing hypothesis for I1 directly; identifying whether this or another axis (parametric robustness, evolvability, upstream-regulator correlations) separates I1 from I3, and testing it in synthetic circuits in which both architectures can be built and stimulated; and replacing the protein-burden proxy with direct fitness measurements such as those of Refs. [27, 29].

## A. Reproducibility

All numerical experiments use `numpy.random.default_rng` with fixed seeds. The parameter scan uses seed 0; the Gillespie replicates use seeds 1, …, *R* within each scan point; the bootstrap analyses use seed 0. The code base is implemented in Python with Numba acceleration for the Gillespie kernel, NumPy/SciPy for the Lyapunov solver and statistical tests, and Matplotlib for figures. The full source code for the analyses presented here, including the configuration files needed to regenerate all main-text and supplementary figures, is available at https://github.com/Rafther0112/FFL_Research [56].

## ACKNOWLEDGMENTS

We thank the Systems Biology Group at Universidad de los Andes for useful discussions.

## Appendix A Derivation of the ten input gate functions

This appendix derives Eqs. (5)–( from a thermo-dynamic occupancy model of the regulated promoter. The basic construction follows Bintu and collaborators [45, 46].

### 1. Thermodynamic framework

Consider a promoter that can be occupied by two transcription factors, *X* and *Y*, each acting either as an activator or as a repressor. Let *g*_X_, *g*_Y_ ∈ [0, 1] denote the active-state occupancies of the two factors, defined via Hill functions as in the main text (Eqs. (2)–( where the sign of the regulation determines whether the activator or repressor form of the Hill function applies. The promoter has a set of microscopic states, one for each combination of bound or unbound factors and of activator or repressor configurations, and to each state we associate a statistical weight and a transcriptional activity. The mean transcription rate of *Z* is then the ensemble average of the activity weighted by the Boltzmann weights of each state.

In our parametrization the dissociation constants for each configuration are absorbed into the Hill constants *K*_XZ_, *K*_Y Z_ that define *g*_X_ and *g*_Y_, leaving the microscopic parameters *K*_1_, …, *K*_5_ and *a* of Ref. [46] as free parameters. For the analytical derivation we set *K*_i_ = 1 for *i* = 1, …, 5 and *a* = 1; this is the simplifying choice that makes the resulting input functions independent of the absolute promoter-TF binding strengths while retaining their structural distinctness. Within this choice, the partition function of the regulated promoter has the form *Z*_stat_(*g*_X_, *g*_Y_ ) = 1 + *g*_X_ + *g*_Y_ + *g*_X_ *g*_Y_ plus auxiliary cross-terms that depend on the configuration of the bound factors, and the mean transcription rate is the weighted sum over configurations.

### 2. Boolean limits: OR and AND

The simplest two configurations are the Boolean OR and AND gates. The OR gate produces nonzero transcription whenever either factor is active; the corresponding mean transcription rate (normalized so that it returns values in [0, 1]) is *g*_X_ + *g*_Y_ − *g*_X_ *g*_Y_, giving Eq. (5) after multiplication by *β*_z_. The AND gate, conversely, requires both factors to be active, yielding the product *g*_X_ *g*_Y_ of Eq. (6).

### 3. Asymmetric activator-repressor logics (L3, L4, L13, L14)

A more general class of two-input regulation arises when one factor acts as an activator and the other as a repressor at the promoter, with binding interactions that are not symmetric between the two. For L3, with *g*_X_ the activator and *g*_Y_ the repressor occupancy, the partition function at the parameter choices above is *Z*_stat_ = (1 + *g*_Y_ )(1 + *g*_X_ + *g*_Y_ ) and the productive-state fraction is *g*_X_ */Z*_stat_, giving Eq. (7). Exchanging the roles of *X* and *Y* gives Eq. (8). Equations (10) and (9) are the complements of Eqs. (7) and (8), respectively, and correspond to the non-productive state fraction.

### 4. Symmetric XOR/XNOR logics (L7, L10)

The XOR-like logic L7 corresponds to the partition function (1 + *g*_X_)(1 + *g*_Y_ ) with a productive-state weight that vanishes when both factors are bound or both are unbound and is nonzero when only one is bound. This gives the numerator *g*_X_ + *g*_Y_ 2*g*_X_ *g*_Y_ of Eq. (11); its complement L10 (XNOR-like) is Eq. (12).

### 5. NAND- and NOR-like Hill complements (L15 and L5)

The remaining two logics are NAND- and NOR-like complements in which each input is passed through an additional Hill function. For L15 (NAND-like) we have

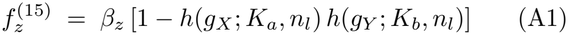

where 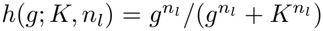 is a secondary Hill. The defaults *K*_a_ = *K*_b_ = 0.5 and *n*_l_ = 2 recover Eq. (13). For L5 (NOR-like, asymmetric in the two factors) we have

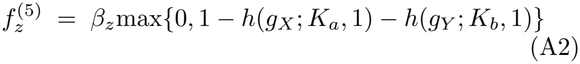

with defaults *K*_a_ = 0.5 and *K*_b_ = 1, recovering Eq. (14). The max operation enforces non-negativity of the transcription rate, which can otherwise be violated by the additive form. The secondary-Hill constants *K*_a_, *K*_b_ and cooperativity *n*_l_ are held at these fixed values rather than sampled, because they parametrize the shape of two specific logic functions rather than a kinetic property of a given circuit; the gate-simplex analysis already averages over the choice of logic, so a distribution over these internal constants would only widen the same average. Because the secondary Hill in Eqs. (13) and (14) is composed on top of the primary occupancies *g*_X_, *g*_Y_, which are themselves Hill functions of coefficient *n*, L5 and L15 operate at an effective cooperativity *nn*_l_ and are therefore steeper than the other eight logics; this steepness is a property of the NAND- and NOR-like forms, not an artifact of the parametrization.

### 6. Closed-form Jacobian entries for AND and OR gates

The Jacobian matrix (16) requires the partial derivatives *∂*_X_ *f*_z_ |_⋆_ and *∂*_Y_ *f*_z_| _⋆_ evaluated at the deterministic steady state. For the two canonical Boolean gates these are obtained in closed form by applying the chain rule to Eqs. (5)–(6).

For the OR gate, 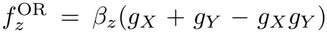, the partial derivatives are

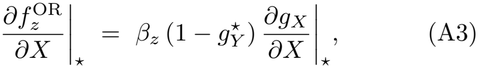

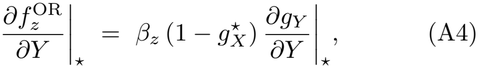

where 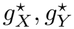 are the steady-state occupancies and

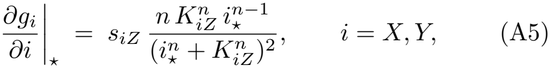

is the steady-state slope of the activator (*s*_iZ_ = +1) or repressor (*s*_iZ_ = − 1) Hill function with respect to its argument.

For the AND gate, 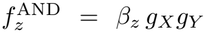, the partial derivatives are simply

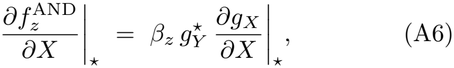

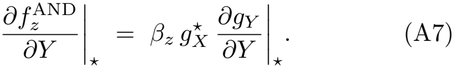

The remaining eight logics have closed-form derivatives of more cumbersome appearance, which we evaluate numerically. Equations (A3)–( together with the analogous expression for *∂*_X_ *f*_y_, fully specify **A**(*E*) for any FFL under AND or OR gating and suffice to reproduce the analytical capacity results of Sec. VI for these two gates.

## Appendix B Numerical methods

### 1. Parameter ranges

The kinetic parameters are sampled log-uniformly within the following ranges: *β*_y_, *β*_z_ ∈ [5, 80]; *K*_XY_, *K*_XZ_ ∈ [150, 2000]; *K*_Y Z_ ∈ [300, 4000]. The Hill coefficient takes values *n* {2, 4 } . The degradation rates are fixed: *γ*_X_ = *γ*_Y_ = *γ*_Z_ = 1 and *µ*_X_ = *µ*_Y_ = *µ*_Z_ = 0.05. The translation rates are fixed: *K*_pX_ = *K*_pY_ = *K*_pZ_ = 5. These are wide absolute ranges rather than ranges tied to the achievable level of each regulator. With the rates above the mean input is ⟨*X*⟩ = *K*_pX_ *K*_X_ */*(*γ*_X_ *µ*_X_) = 100 *K*_X_, so over *K*_X_∈ [0.5, 10] the input spans roughly 50 to 1000 molecules, and dissociation constants sampled up to 2000 can exceed the range a regulator reaches. A fraction of the sampled circuits therefore never cross their own dissociation constants and respond weakly to the environment. We use absolute ranges deliberately, so as not to impose responsiveness a priori: restricting each *K* to the achievable range of its regulator would build functional circuits into the prior by construction, whereas the wide prior lets the cost-information analysis, rather than the sampling, decide which architectures are favored. We verify in Sec. VI C, with two filters that remove the non-responsive samples while preserving the paired comparison, that the rankings and the near-equivalence of I1 and I3 are not artifacts of the wide prior.

### 2. Lyapunov solver

The Lyapunov equation (18) is solved numerically using the `scipy.linalg.solve continuous lyapunov` routine. The local gain *g*(*E*) = *d* ⟨*Z*⟩ */dE* is computed by central differences on a logarithmically spaced grid of *K*_X_ values 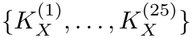 spanning [0.5, 10]. The Brunel-Nadal-Laughlin integral of Eq. (22) is evaluated by trapezoidal quadrature on the same grid.

### 3. Gillespie simulations

Stochastic simulations of the master equation are performed with the standard Gillespie algorithm [38] implemented in Numba-accelerated Python. Steady-state analyses use a burn-in of 200 time units, followed by sampling over 1000 time units; transient analyses use a 200– unit pre-equilibration at *K*_X_ = 1, followed by a step to each of the eight values 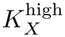 of Sec. VI D and tracking of *Z*(*t*) over *T* = 300 time units, sampled every 4 time units.

### 4. Mutual information estimator

The mutual information *Î*(*E*; *Z*) is estimated with the plug-in estimator with Miller-Madow bias correction [39], using 8 quantile bins over the empirical distribution of *Z* and a discrete prior of 8 log-spaced *K*_X_ values in [1, 8]. The analytical capacity, which involves no bins or replicates, agrees with this estimator in correlation and in within-family ranking (Sec. IV D and App. C 1), and for I1 and I3 under (OR, *n* = 2) a finer plug-in estimator and a binning-free KSG estimator give the same information values (Sec. IV).

### 5. Cross-validation between Lyapunov and SSA

For each of the *N* = 1000 log-uniform parameter samples in the stationary stratified Pareto analysis we recompute the analytical Lyapunov channel capacity (Eq. (22)) under identical parameters and compare it pointwise against the empirical *Î*_SSA_ at the same sample. Pearson and Spearman correlations are computed across all paired measurements within each (family, gate, *n*) sub-group.

## Appendix C Additional analyses

### 1. Within-family rank correlations between Lyapunov and SSA

Table II lists, for each (family, gate, *n*) subgroup, the within-family ranking of variants by mean information under the two estimators. The transpositions in three incoherent subgroups involve different pairs, in each case the two variants closest in mean information, as expected when two estimators are applied to tightly spaced values. The table documents the agreement between the estimators; it is not a cost-information ranking.

### 2. Bootstrap confidence intervals and paired I1/I3 test

Table III reports the bootstrap 95% confidence intervals on the stratified Pareto shares of Fig. 3, together with the result of the Wilcoxon signed-rank paired test on the per-sample difference *D*_k_ = share(I1)_k_ − share(I3)_k_.

### 3. Per-subgroup offset between Lyapunov and SSA

Table IV gives the linear fit of *C* against *Î*_SSA_ in each subgroup, with 95% bootstrap intervals over 2000 resamples, to test whether the offset of Sec. IV D is larger for the non-monotonic incoherent variants. The underlying paired values are shown in Fig. 6.

**FIG 6.**
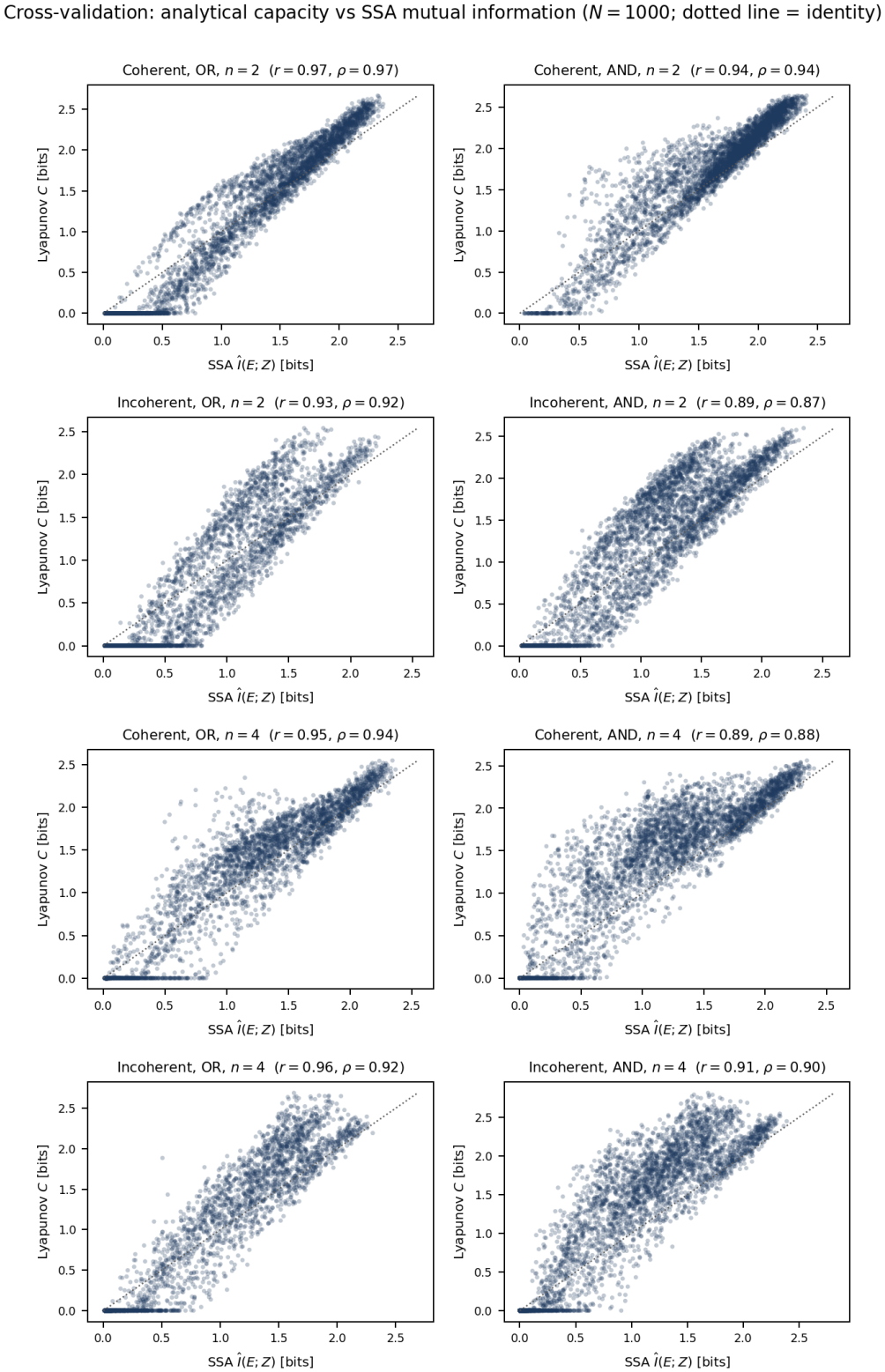
Cross-validation between the analytical capacity *C* and the Gillespie estimate *Î*_SSA_(*E*; *Z*) at *N* = 1000 log-uniform parameter samples. Panels show the eight (family, gate, *n*) subgroups in row-major order: coherent OR and AND at *n* = 2; incoherent OR and AND at *n* = 2; coherent OR and AND at *n* = 4; incoherent OR and AND at *n* = 4. Within each panel the four variants of the family are overlaid; Pearson and Spearman correlations are printed in each title, and the dotted line is the identity. The Pearson correlations lie in [0.80, 0.95] across subgroups.

The slope means are 1.02 for the coherent and 1.12 for the incoherent family, and a Mann-Whitney test over the four subgroups per family finds no difference (*p* = 0.49). The coherent slopes straddle unity (from 0.82 to 1.23), so there is no systematic family-dependent offset; the aggregate excess of the capacity over the SSA estimate (Sec. IV D) reflects the optimal prior and the wider integration domain, not a branch-degeneracy term. The clipped fraction, by contrast, is markedly higher in the incoherent OR subgroups (about 45%) than in the coherent AND ones (as low as 3%), confirming that the small-noise floor is reached where the non-monotonic response is least informative (Sec. VI C).

### 4. Per-gate Pareto shares

Figure 7 resolves the stationary Pareto shares of Fig. 3 across all ten gate logics of Sec. III B, at *n* = 2 and 4 and for both families. C1 leads the coherent family under six of the ten gates but is displaced by C3 under the NAND- and NOR-like logics (L15, L5), the XOR-like logic L7, and the XNOR-like logic L13. Among incoherent variants the lead is split evenly: I3 leads under OR, AND, L4, L10, and L14, and I1 under L3, L5, L7, L13, and L15, five gates each at both *n*, with no variant leading under all gates. The gate therefore determines whether I1 or I3 is ahead, with differences as large as +43 (I1 over I3 under L13) and − 45 percentage points (under L14). These per-gate shares are the *P*_*l*_(FFL) that enter the gate-simplex analysis of Sec. VI F.

**FIG 7.**
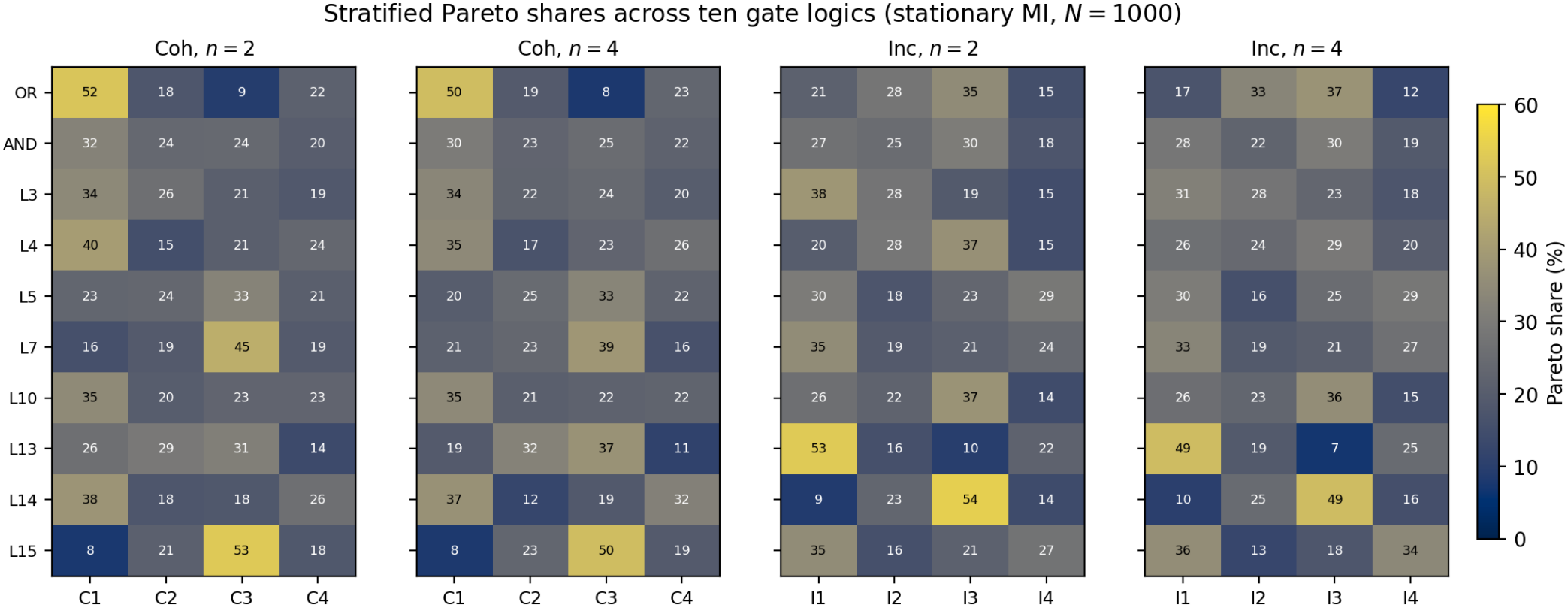
Stratified Pareto shares (%) across all ten input gate logics (rows) for the coherent and incoherent families at Hill coefficients *n* = 2 and *n* = 4 (column groups), computed from *N* = 1000 log-uniform parameter samples and shown as a cividis map (dark low, bright high). Each column within a panel is one within-family variant competing on the Pareto frontier; shares are computed with the same 1*/k* credit rule as Fig. 3 and sum to 100% per panel. The two canonical gates OR and AND are the first two rows; the remaining eight are the additional logics L3, L4, L5, L7, L10, L13, L14, L15 defined in Appendix A. The bright cells trace which variant leads under each logic: C1 under most gates, C3 under the NAND–, NOR–, XOR– and XNOR-like logics, and I1 and I3 alternating across the incoherent gates.

### 5. Threshold sensitivity of the stationary results

Figure 8 shows the common-support test of Sec. VI C: the mean per-sample difference *D*_k_ between I1 and I3, with its bootstrap interval, and the number of retained samples, as the threshold above which all four incoherent variants must be informative is swept from 0 to 0.5 bits. The difference never crosses to favor I1; under OR the strict criterion exhausts the sample, widening the interval through loss of power.

**FIG 8.**
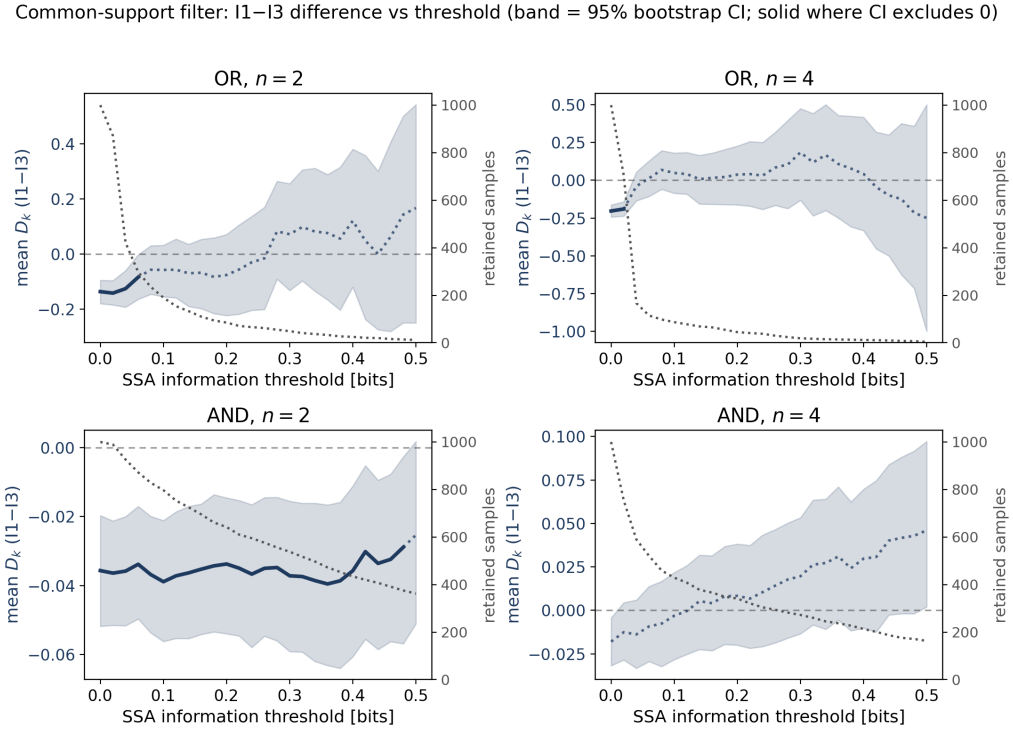
Common-support filter for the two canonical gates. Each panel shows the mean per-sample difference *D*_*k*_ = share(I1) − share(I3) (navy, left axis) with its 95% bootstrap interval (shaded band), and the number of retained samples (grey dotted, right axis), against the SSA information thresh-old above which all four incoherent variants must lie for a sample to be kept. Under AND the filter retains 640–900 samples and the interval stays on the *D*_*k*_ *<* 0 side (or relaxes to zero at *n* = 4) without ever crossing to favor I1. Under OR the strict criterion collapses the retained sample (I2 and I3 are the variants most often unresponsive), so the interval widens through loss of power, not a change of sign. No panel shows a reversal toward I1.

### 6. Empirical census counts

Table V lists the FFL counts of Ref. [21] next to the stationary Pareto shares at *n* = 2 under OR gating, for the two organisms with well-characterized censuses.

